# JCAD couples tight junction condensates to actin and RhoA to maintain the endothelial barrier

**DOI:** 10.64898/2026.05.19.725802

**Authors:** Kyle A Jacobs, Yoon-Gu Jang, Fang-Shiuan Leung, Lakyn N Mayo, Torsten Wittmann, Jeffrey O Bush, Matthew L Kutys

**Affiliations:** Department of Cell and Tissue Biology, University of California San Francisco, San Francisco, CA; Cardiovascular Research Institute, University of California San Francisco, San Francisco, CA; Biomedical Sciences Graduate Program, University of California San Francisco, San Francisco, CA; Program in Craniofacial Biology, University of California San Francisco, San Francisco, CA; Institute for Human Genetics, University of California San Francisco, San Francisco, CA; Medical Scientist Training Program, University of California San Francisco, San Francisco, CA

**Author notes:** Corresponding author: Matthew L Kutys, PhD 513 Parnassus Avenue, Box 0512, HSW 618, San Francisco, CA 94143.

## Abstract

How endothelial cell-cell junctions integrate cytoskeletal, adhesive, and local signaling networks to maintain vascular barrier integrity remains incompletely defined. Here, we identify junctional cadherin 5-associated protein (JCAD) as a modular scaffold that organizes endothelial tight junction architecture by coupling junctional condensates to actin and RhoA signaling. Genetic deletion of *Jcad* in mice does not affect baseline vascular permeability but causes inflammation-dependent barrier hyperpermeability. JCAD depletion in primary human endothelial cells disrupts tight junction continuity and increases paracellular permeability. Mechanistically, JCAD localizes to ZO-1-positive tight junctions independently of VE-cadherin, directly binds filamentous actin, and forms dynamic actin-associated condensates at cell-cell contacts. Structure-function analysis reveals separable domains mediating tight junction targeting and actin binding, establishing a bipartite architecture that distinctly coordinates junctional signaling and cytoskeletal coupling. Together, these findings identify JCAD as a cell-cell adhesion scaffold that integrates the phase-separated tight junction plaque with actin and RhoA-dependent mechanics, enabling endothelial barrier adaptation to inflammatory stress.

## Introduction

Endothelial cells line blood vessel lumens and establish a selectively permeable barrier between blood and the extravascular space. This barrier must be dynamically regulated – transiently loosening to permit leukocyte extravasation during immune surveillance and inflammation (Muller, 2003), while rapidly recovering to prevent sustained leakage. Dysfunction of this regulation contributes to the pathogenesis of diverse diseases, including atherosclerosis (Libby, 2002; Ross and Glomset, 1976; Gimbrone and García-Cardeña, 2016), edema (Corada et al., 1999; Senger et al., 1983), and vascular dementias (Sweeney et al., 2018; Montagne et al., 2017). Despite the importance of endothelial barrier control in vascular disease, the molecular mechanisms coordinating cell-cell junction remodeling remain incompletely defined, limiting the development of strategies to therapeutically control vascular permeability.

Endothelial barrier function is primarily mediated by intercellular junctions, including adherens junctions (AJs) and tight junctions (TJs). AJs are organized around the transmembrane adhesion receptor VE-cadherin and its associated intracellular catenin complex, while TJs are composed of tetraspan transmembrane proteins (claudins/occludins) and their associated intracellular zonula occludens (ZO) scaffold proteins. Beyond providing structural cohesion, these intracellular junctional plaques act as signaling nexuses that regulate numerous cellular processes including mechanotransduction (Conway et al., 2013; Barry et al., 2015), proliferation (Grazia Lampugnani et al., 2003; Lampugnani et al., 2006), and transcriptional programs (Giampietro et al., 2015; Morini et al., 2018; Taddei et al., 2008).

How AJ and TJ plaques coordinate adhesion remodeling in response to physiological and pathological stimuli remains unclear. TJ assembly is driven in part by phase separation of ZO proteins into condensates that concentrate actin, promoting filament polymerization and plaque growth (Beutel et al., 2019; Schwayer et al., 2019; Pombo-García et al., 2024; Sun et al., 2025). AJs are similarly regulated by junction-associated actin: branched actin networks polymerize at the plasma membrane to physically push cell-cell interfaces together (Efimova and Svitkina, 2018), while cortical actomyosin bundles simultaneously couple to the adhesion plaque to apply tension and strengthen adhesions through an α-catenin-mediated catch-bond mechanism (Xu et al., 2020; Heuzé et al., 2019). Reciprocally, TJ and AJ plaques scaffold key regulators of RhoA signaling, including GEF-H1 (Citi et al., 2009) and p190RhoGAP (Wildenberg et al., 2006), respectively, enabling local control of actin dynamics and barrier function. In the endothelium, this spatially restricted Rho GTPase signaling is critical for transient permeability responses to inflammation (Eckenstaler et al., 2022). However, the molecular scaffolds that integrate junctional organization with actin and Rho GTPase signaling remain incompletely defined.

We recently reported junctional cadherin 5 associated protein (JCAD; also known as junctional protein associated with coronary artery disease) as a highly enriched protein proximal to the VE-cadherin intracellular domain in a proximity biotinylation proteomic screen (Mayo et al., 2026; Xia et al., 2025). JCAD is a cell-cell adhesion-associated protein expressed broadly, including in the vascular endothelium (Akashi et al., 2011), the trabecular mesh (Maddala and Rao, 2020), and the perineurium (Hiraoka et al., 2023). JCAD loss has been associated with impaired Hippo and PI3K/AKT signaling (Jones et al., 2018; Ye et al., 2017; Liberale et al., 2023), and JCAD-deficient mice exhibit impaired tumor xenograft angiogenesis (Hara et al., 2017; Ye et al., 2017) and decreased atherosclerotic burden on a high fat diet (Douglas et al., 2020; Xu et al., 2019). Despite these prior studies, whether JCAD has a direct role in organizing or regulating the cell-cell adhesion plaque remains unknown.

Here, we identify JCAD as a cell-cell adhesion scaffold protein that localizes to TJs independent of VE-cadherin, directly binds actin, and forms dynamic, actin-associated condensates at endothelial cell-cell contacts. Using complementary *in vivo* and *in vitro* models, we establish that JCAD is crucial to maintain junctional integrity during inflammatory stress. We define JCAD domains that mediate TJ targeting and actin association, defining a modular architecture by which JCAD achieves selective localization to distinct junctional compartments. Finally, we find that JCAD regulates endothelial responses to thrombin and basal actin dynamics, and selectively binds GTP-bound RhoA, positioning it to locally coordinate RhoA-dependent actomyosin dynamics at the TJ plaque.

Together, these findings identify JCAD as a previously unrecognized component of the endothelial cell-cell adhesion machinery that links TJ condensates to cytoskeletal signaling, revealing how endothelial barriers integrate mechanical and inflammatory cues.

## Results

### JCAD is spatially proximal to the VE-cadherin intracellular domain and regulates vascular barrier function in mice

To identify uncharacterized proteins organizing the endothelial junctional plaque, we conducted a proximity-dependent biotinylation (BioID) screen in primary human microvascular endothelial cells (hMVECs) by tagging the VE-cadherin intracellular domain with the promiscuous biotin ligase BirA*, enabling mass spectrometric identification of proteins residing within the ∼10 nm labeling radius (Figure 1A) (Kim et al., 2014; Xia et al., 2025; Mayo et al., 2026). Among the enriched proximal proteins, we identified JCAD, a ∼145 kDa protein first described as a coronary artery disease risk locus (Peden et al., 2011; Erdmann et al., 2011), with few known functions. We validated JCAD biotinylation by streptavidin affinity precipitation of biotinylated proteins and immunoblotting. JCAD was strongly enriched in the VE-cadherin-BirA* streptavidin precipitation compared to the wildtype control (Figure 1B), confirming its spatial proximity to the VE-cadherin intracellular domain.

**Figure 1:**
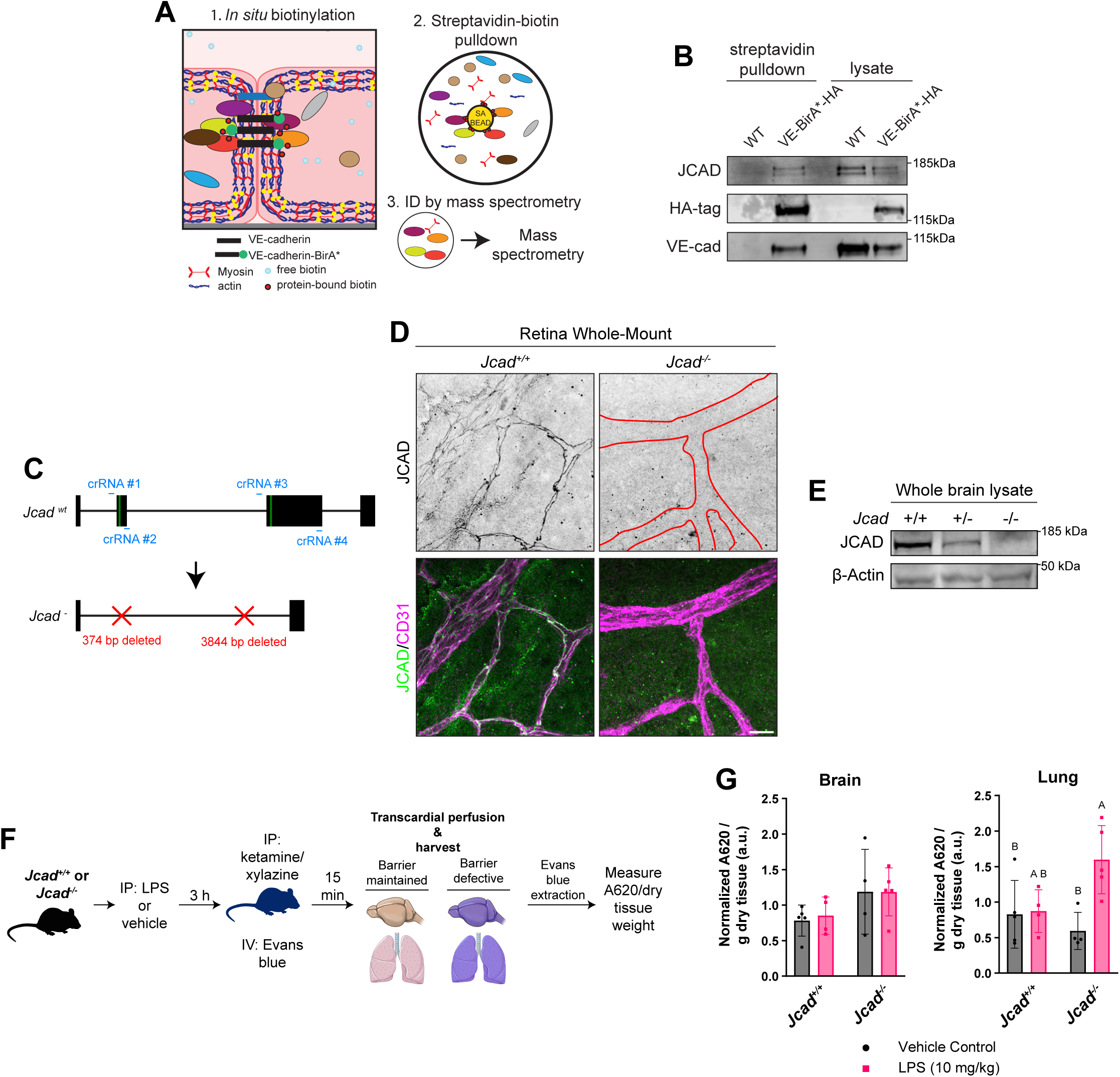
JCAD is a VE-cadherin proximal protein required for maintenance of vascular barrier function in mice. **(A)** Schematic of VE-cadherin proximity-dependent biotinylation screen. **(B)** Western blot of affinity purified biotinylated proteins from wildtype and VE-cadherin-BirA* expressing hMVECs. The HA and VE-cadherin panels of this blot were previously published in (Mayo et al., 2026). **(C)** Schematic of *Jcad* locus exon organization, crRNA target sites, and CRISPR-mediated deletions. Green bands, positions of in-frame start codons. **(D)** Immunostaining for JCAD and CD31 in whole-mount retinas from *Jcad^+/+^* and *Jcad^-/-^* mice. Scale bar, 15µm. **(E)** Western blot demonstrating JCAD protein loss in whole brain lysates from *Jcad^+/+^*, *Jcad^+/-^*, and *Jcad^-/-^* mice. **(F)** Schematic of *in vivo* vascular barrier assay. **(G)** Quantification of Evans blue dye extraction. Values of absorbance at 620 nm per gram of dry lung/brain tissue were normalized to the mean of each litter. Groups not sharing a letter are significantly different (two-way ANOVA with Holm-Šídák’s post hoc test; α = 0.05). N = 4 independent litters (male and female; ages P21-P75); each point represents one animal.

Cell-cell adhesion integrity is critical for vascular endothelial barrier function. To test whether JCAD plays a role in maintaining barrier integrity, we generated JCAD-deficient mice by deleting *Jcad* exons containing in-frame start codons (Figure 1C; Figure S1A) using the CRISPR/Cas9-based approach improved-Genome editing via Oviductal Nucleic Acids Delivery (i-GONAD) (Gurumurthy et al., 2019; Ohtsuka et al., 2018). Immunoblot analysis of whole-brain lysates and immunostaining of whole-mount retinas confirmed loss of JCAD protein in *Jcad* deficient (*Jcad^-/-^*) mice (Figure 1D-E). *Jcad^-/-^* mice were born at expected Mendelian ratios (Figure S1B-C) and exhibited no overt developmental defects.

To determine if *Jcad^-/-^* mice exhibit defects in vascular barrier regulation, we conducted an Evans blue vascular extravasation assay (Radu and Chernoff, 2013). When injected intravenously, Evans blue binds circulating albumin (∼70 kDa), thereby serving as a tracer for plasma protein extravasation (Lindner and Heinle, 1982). In addition to testing extravasation under control conditions, we also challenged mice with acute lipopolysaccharide (LPS) exposure, a model of endotoxemia-induced barrier disruption, to determine whether JCAD is required to maintain barrier integrity at baseline and under inflammatory conditions (Figure 1F). Three hours after LPS administration, Evans blue extravasation in the brain and lung – two vascular beds with fundamentally distinct barrier properties – was assessed. Brain vasculature enforces a highly restrictive blood-brain barrier, while the pulmonary vasculature maintains a more dynamically regulated barrier that is acutely sensitive to inflammatory insult.

In the brain, two-way ANOVA revealed no significant main effects or interactions, although genotype approached significance (p = 0.0546) (Figure 1G). In contrast, in the lung, there was a significant genotype × LPS interaction (p = 0.0262) and a significant main effect of LPS (p = 0.0164), while genotype alone was not significant (p = 0.2239). Post-hoc analyses demonstrated that *Jcad^-/-^* mice treated with LPS were significantly different from both *Jcad^+/+^* and *Jcad^-/-^* mice treated with vehicle (p = 0.0469 and p = 0.0146, respectively; Figure 1G), with a near-significant difference between *Jcad^+/+^* and *Jcad^-/-^* treated with LPS (p = 0.0730), consistent with the interaction effect. Together, these data indicate that JCAD is dispensable for baseline vascular integrity but is required to maintain barrier function during inflammatory stress.

### JCAD maintains hMVEC tight junction integrity and endothelial barrier function

Because JCAD-deficient mice exhibited inflammation-dependent vascular hyperpermeability, we next determined how JCAD contributes to the dynamic regulation of vascular barrier integrity. Using lentivirus-mediated short hairpin RNA (shRNA) (Rubinson et al., 2003), we silenced *JCAD* expression in hMVECs, which reduced JCAD protein levels greater than 10-fold (Figure 2A-B). JCAD robustly localized to cell-cell junctions by immunofluorescence in control hMVECs expressing a non-targeting (Control) shRNA but was absent in JCAD knockdown hMVECs (JCAD KD; Figure 2C).

**Figure 2:**
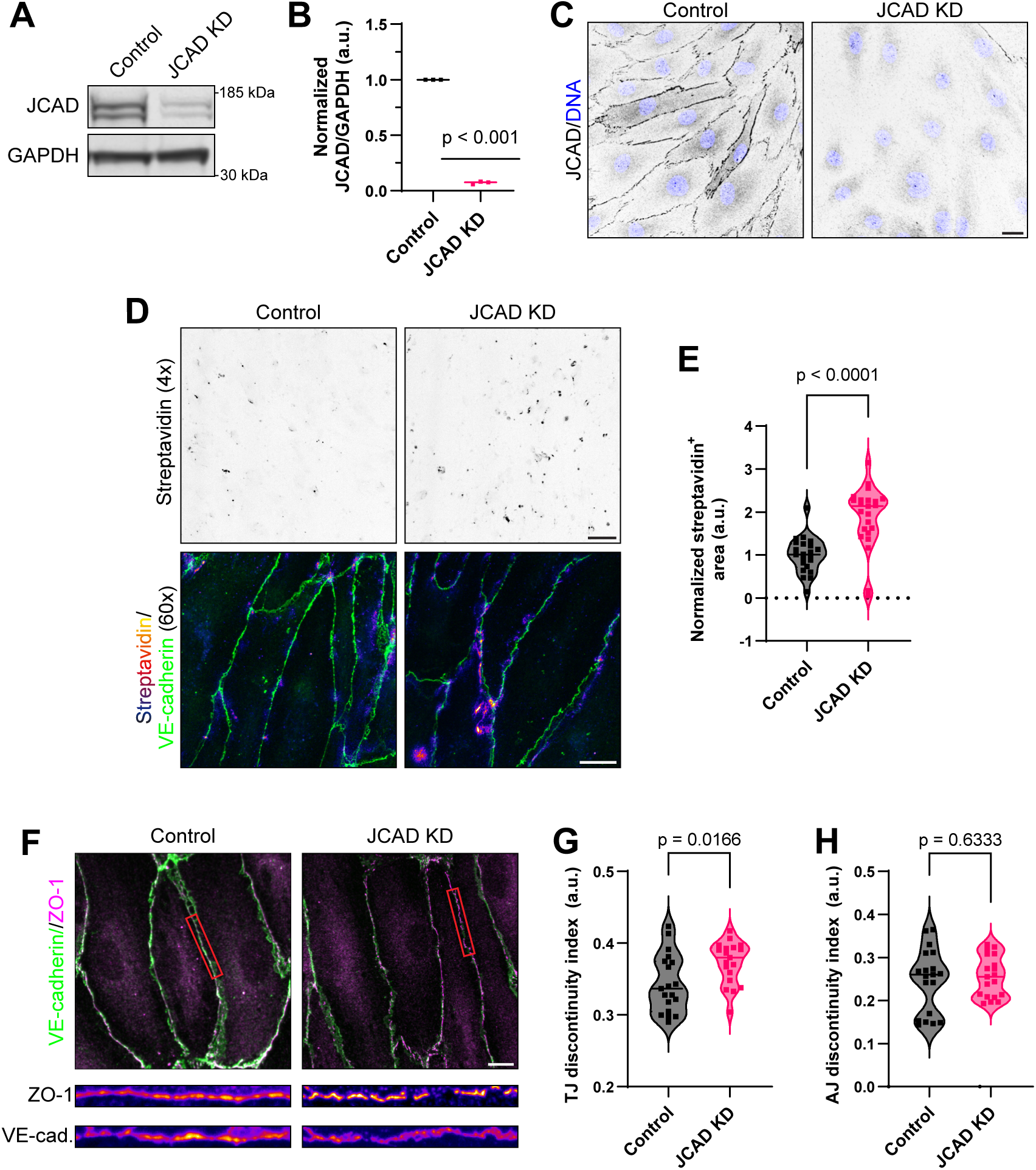
JCAD regulates barrier permeability and tight junction integrity in human microvascular endothelial cells. **(A)** Western blot of Control and JCAD KD hMVECs immunoblotted for JCAD and GAPDH. **(B)** Quantification of JCAD KD western blot validation across multiple replicates (N = 3 independent experiments). **(C)** Immunostaining for JCAD in Control and JCAD KD hMVECs. Scale bar, 20 µm. **(D)** Representative streptavidin staining in Control and JCAD KD cells plated on biotinylated fibronectin-coated coverslips. Low magnification (top; scale bar, 200 µm) shows labeling across a wide field, while high magnification (bottom; scale bar, 15 µm) reveals increased streptavidin intensity at cell-cell boundaries marked by VE-cadherin immunostaining (green). **(E)** Quantification of normalized streptavidin-labeled area across multiple replicates (N = 3 independent experiments, n = 6-8 fields of view per N; Mann-Whitney test). **(F)** Representative images of VE-cadherin and ZO-1 immunostaining in Control and JCAD KD hMVECs. Red boxes indicate the regions shown in the inset below (intensity-scaled heatmap). Scale bar, 15 µm. **(G/H)** Quantification of tight and adherens junction discontinuity indices over multiple independent replicates (N = 3 independent experiments, n = 5-9 fields of view per N, Welch’s *t*-test).

To assay the effect of JCAD KD on endothelial barrier integrity, we conducted an *in vitro* barrier function assay (Aw et al., 2025; Rathod et al., 2024). Control and JCAD KD hMVECs were seeded onto coverslips coated with biotinylated fibronectin and laminar fluid shear stress was applied to promote junction maturation and barrier formation (Hahn and Schwartz, 2009). Immediately prior to fixation, coverslips were incubated with fluorescently labeled streptavidin for two minutes to assay the permeability of the hMVEC monolayer. Fluorescence microscopy revealed that JCAD KD cells exhibited a significantly greater streptavidin-labeled area relative to Control cells (Figure 2D-E; p < 0.0001). High-magnification imaging revealed that streptavidin labeling of the substrate correlated with the location of hMVEC cell-cell interfaces, labeled by VE-cadherin immunostaining (Figure 2D, bottom), indicating that the increased monolayer permeability is likely caused by increased paracellular solute transit.

To determine whether JCAD KD-induced barrier dysfunction stems from impaired cell-cell junctional integrity, we immunostained for ZO-1 and VE-cadherin in JCAD KD and Control cells. Cells were seeded on compliant 2.5 kPa polyacrylamide gel coated with globular collagen I and subjected to laminar fluid shear stress. Quantitative analysis of ZO-1 immunostaining at cell-cell interfaces showed that JCAD KD cells had significantly disrupted TJs, reflected by a higher junctional discontinuity index, defined as the length-weighted mean coefficient of variation of junctional fluorescence intensity, compared to Control cells (Figure 2F-G; p = 0.0166). In contrast, the VE-cadherin discontinuity index was not significantly different between JCAD KD and control cells (Figure 2H; p = 0.6333), indicating that AJ integrity was not significantly disrupted. Together, these findings demonstrate that JCAD loss selectively disrupts TJ continuity without significantly affecting AJ continuity, identifying JCAD as a novel regulator of TJ architecture and endothelial barrier integrity.

### JCAD localizes to tight junctions independently of VE-cadherin

Having established that JCAD regulates endothelial barrier function both in mice and in hMVECs in culture, we next defined how JCAD localizes to the endothelial cell-cell adhesion complex to better understand the molecular mechanism by which it contributes to junction integrity.

JCAD was originally described as an AJ-associated protein (Akashi et al., 2011), consistent with its identification in spatial proximity to the VE-cadherin intracellular domain (Figure 1B). However, in high-resolution confocal images of hMVECs co-immunostained for JCAD, VE-cadherin, and ZO-1, JCAD localization was not consistent with classical endothelial AJ architecture. JCAD localized in a tight, continuous, linear pattern along cell-cell interfaces, closely mirroring ZO-1 (Figure 3A). Notably, JCAD was excluded from the junctional reticulations characteristic of VE-cadherin AJs (Figure 3Ai-ii, arrows) (Cao and Schnittler, 2019). Quantitative analysis of fluorescent signal correlation across junctional line scans confirmed significantly higher JCAD-ZO-1 correlation compared to JCAD-VE-cadherin correlation (Figure 3B; p = 0.0003).

**Figure 3:**
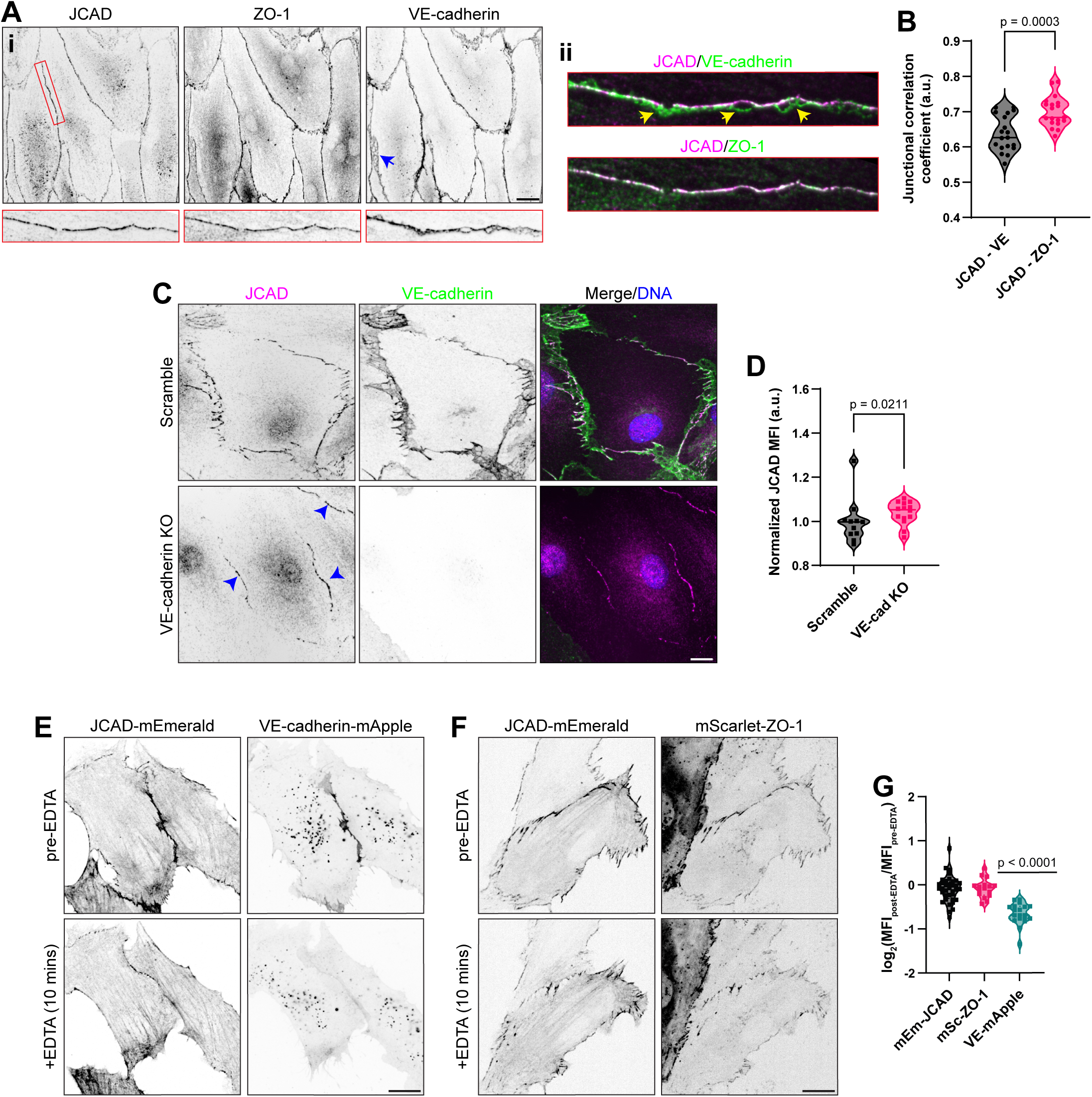
JCAD localizes to cell-cell interfaces independent of VE-cadherin. **(A) (i)** Immunostaining for JCAD, VE-cadherin, and ZO-1 in hMVECs. Red box, location of inset. Blue arrow, reticulated adherens junction lacking JCAD staining. Scale bar, 20 µm. **(ii)** Composite images demonstrating JCAD-VE-cadherin and JCAD-ZO-1 overlap. Yellow arrows, regions with no JCAD-VE-cadherin overlap. **(B)** Quantification of JCAD-VE-cadherin and JCAD-ZO-1 correlation coefficients at cell-cell adhesions (N = 3 independent experiments, n = 6-8 fields of view per N, Welch’s t test). **(C)** Immunostaining for JCAD and VE-cadherin in Scramble and VE-cadherin knockout (KO) hMVECs. Scale bar, 20 µm. Arrowheads, JCAD localization at junction without VE-cadherin. **(D)** Quantification of JCAD normalized mean fluorescence intensity at cell-cell interfaces (N = 3 independent experiments, n = 3-5 fields of view per N, Mann-Whitney test). **(E/F)** Live-cell imaging of hMVECs expressing JCAD-mEmerald and either (E) VE-cadherin-mApple or (F) mScarlet-ZO-1 before and after treatment with EDTA (2.5 mM, 10 min). Scale bars, 15 µm. **(G)** Quantification of log_2_ of the fold change in mean junctional fluorescence intensity of each protein before and after EDTA chelation (N = 3 independent experiments, n = 5-8 fields of view per N, one sample *t* test).

To confirm that this localization pattern is not specific to dermal hMVECs, we immunostained dermal lymphatic endothelial cells and blood endothelial cells of differing organotypic origins including brain microvascular, coronary artery, and umbilical vein. In all cases, JCAD consistently showed significant overlap with ZO-1 in a linear, non-reticulating pattern (Figure S2A). Collectively, these data suggested that while JCAD resides within nanoscale proximity to the VE-cadherin intracellular domain, it may not be an AJ-associated protein as previously classified, prompting us to investigate the mechanism of JCAD junctional targeting.

We next asked whether genetic ablation of AJs by VE-cadherin depletion results in loss of JCAD junctional localization. We utilized previously generated and validated VE-cadherin CRISPR/Cas9 knockout (KO) hMVECs (Polacheck et al., 2017; Xia et al., 2025). Immunostaining for JCAD in VE-cadherin KO hMVECs revealed no decrease in JCAD localization at cell-cell interfaces, and instead, quantification of JCAD mean fluorescence intensity at cell-cell interfaces revealed a modest, yet significant, increase in VE-cadherin KO relative to Scramble (Figure 3C-D; p = 0.0211). Together these data demonstrate that JCAD localizes to junctions in the absence of VE-cadherin.

To further validate this finding in a system lacking VE-cadherin expression entirely, we immunostained for JCAD in non-endothelial cell types that lack VE-cadherin expression (Lampugnani et al., 1992), including U-2 OS osteosarcoma cells and Caco-2 epithelial cells. In both cell lines, JCAD was expressed and localized to a subset of ZO-1-positive TJs (Figure S2B). These findings collectively indicate that JCAD localization to cell-cell interfaces does not require VE-cadherin expression.

To preclude the possibility that JCAD retention at junctions in VE-cadherin KO cells reflects reliance on other cadherins, we performed live-cell imaging during acute AJ disruption via Ca^2+^ chelation (Takeichi, 1977; Simons, 2023). hMVECs co-expressing mEmerald-tagged JCAD together with either VE-cadherin-mApple, to visualize AJs, or mScarlet-ZO-1, to visualize TJs, were imaged before and after EDTA-mediated chelation. At 10 minutes of chelation, we observed significant dissipation of VE-cadherin from cell-cell contacts (Figures 3E & 3F; p < 0.0001), while mScarlet-ZO-1 was unchanged, indicating that at this timepoint AJs have disassembled but TJ contacts persist. mEmerald-JCAD fluorescence intensity at cell-cell interfaces was similarly unchanged at this timepoint (Figure 3G). Together, these data confirm that acute AJ disruption does not displace JCAD from cell-cell contacts, challenging its classification as an AJ protein and instead supporting its localization within the tight junction plaque.

### JCAD directly links tight junctions to the actin cytoskeleton

In live-cell microscopy images of mEmerald-JCAD expressing cells, we identified a subset of cells in which JCAD localized to a filamentous network across the body of the cell in addition to cell-cell contacts (Figure 4A; Video 1). To confirm that this localization pattern is not an artifact of the tagging strategy, we generated multiple independently tagged constructs, including C-terminal JCAD-mEmerald, N- and C-terminal HA-tagged JCAD, and N-terminal mEos2-JCAD. All constructs demonstrated a similar filamentous localization pattern (Figure S3), indicating that this distribution was not tag-dependent.

**Figure 4:**
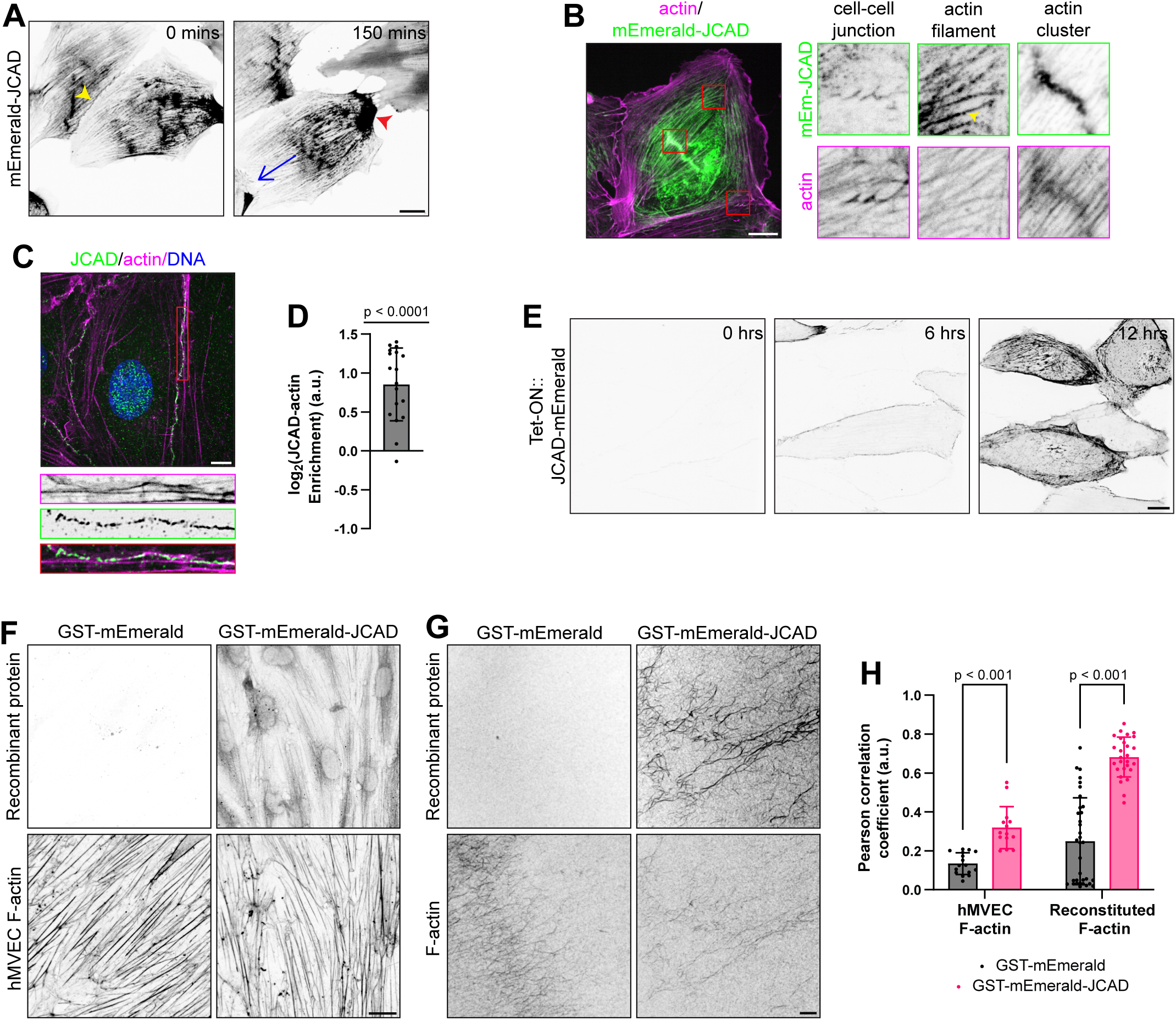
JCAD is a novel actin-binding protein. **(A)** Live-cell imaging of hMVECs expressing mEmerald-JCAD under human cytomegalovirus promoter. Yellow arrowhead, localization to cell-cell interface. Blue arrow, direction of migration. Red arrowhead, trailing edge accumulation. Scale bar, 20 µm. **(B)** Representative image of phalloidin-labeled actin in hMVEC expressing mEmerald-JCAD. mEmerald-JCAD image was log transformed to facilitate visualization of structures across wide intensity range. Red boxes, location of insets. Yellow arrowhead, examples of JCAD punctate localization along actin fibers. Scale bar, 15 µm. **(C)** Immunostaining for JCAD and phalloidin labeling of F-actin in hMVECs. Red box, junctional inset location. Scale bar, 5 µm. **(D)** Quantification of the fraction of JCAD signal overlapping junctional actin expressed as the log₂ ratio of observed overlap to randomized overlap (N = 3 independent experiments, n = 5-9 junctions per experiment; One sample *t* test). **(E)** Representative live-cell images of hMVECs with doxycycline-induced JCAD-mEmerald (TetON::JCAD-mEmerald) expression. Scale bar, 15 µm. **(F)** Representative image of hMVEC fixed and stained with GST-mEmerald or GST-mEmerald-JCAD, followed by phalloidin labeling of F-actin. Scale bar, 15 µm. **(G)** Representative image of *in vitro* reconstituted actin filaments stained with GST-mEmerald or GST-mEmerald-JCAD, followed by phalloidin labeling of F-actin. Scale bar, 5 µm. **(H)** Quantification of correlation between phalloidin labeling of F-actin and mEmerald signal in hMVECs (F) and in reconstituted F-actin (G) (N = 3-4 independent experiments for both; n = 4-13 fields of view per N; Welch’s *t* test).

The polarization of the filamentous network at the trailing edge of migrating hMVECs resembled actomyosin polarization in a migrating cell (Figure 4A, red arrowhead; Video 1) (Vicente-Manzanares et al., 2009; Weißenbruch and Mayor, 2024). We stained for filamentous actin (F-actin) in mEmerald-JCAD expressing cells and observed JCAD localization to a subset of the F-actin network, both at cell-cell interfaces and across the cell body (Figure 4B). Notably, JCAD did not fully overlap individual fibers, but rather appeared as discrete puncta along the filament, with significant accumulation at actin clusters (Figure 4B, inset). Immunostaining for α-tubulin and vimentin in JCAD-mEmerald-expressing cells demonstrated no appreciable overlap with JCAD (Figure S4), indicating that JCAD selectively associates with the actin cytoskeleton among major cytoskeletal networks. Together, these data demonstrate that overexpressed JCAD localized to a subset of the actin cytoskeleton in a punctate pattern.

We next asked whether similar localization occurs at endogenously expressed levels. Endothelial cell-cell adhesions are characterized by junction-proximal actomyosin cables that interface with components of the cell-cell adhesion plaque (Rouaud et al., 2023; Efimova and Svitkina, 2018; Heuzé et al., 2019). To assess whether endogenously expressed JCAD similarly associates with junction-associated actin cables, we performed high-resolution confocal imaging of hMVECs co-stained for JCAD and F-actin. JCAD signal strongly associated with the actin filaments at the cell-cell interface (Figure 4C), and fluorescence intensity-based isolation of junctional F-actin and JCAD signal revealed significant correlation (Figure 4D; p < 0.0001).

Cells exhibiting prominent filamentous localization of JCAD appeared to express it at higher levels than those with only junctional localization. To determine whether higher expression levels of JCAD correlate with the shift of some JCAD from junctional to filamentous localization, we drove JCAD-mEmerald expression in hMVECs using a doxycycline-inducible promoter (Tet-On::JCAD-mEmerald). Time-course live-cell imaging after doxycycline administration revealed a sequential localization pattern. At lower expression levels, JCAD was localized predominantly to cell-cell interfaces. As expression increased, JCAD progressively accumulated along the filamentous network spanning the cell body (Figure 4E; Video 2).

Having demonstrated JCAD co-localization with junctional actin fibers, we next asked whether this interaction is direct or mediated by an adaptor protein. We purified recombinant GST-tagged mEmerald and mEmerald-JCAD from *E. coli* and confirmed full-length JCAD purification by immunoblot analysis (Figure S5). Using these recombinant proteins to stain fixed, permeabilized hMVECs, mEmerald-JCAD – but not mEmerald alone – showed significant overlap with phalloidin-labeled F-actin (Figure 4F), with significantly higher signal correlation than the mEmerald control (Figure 4H; p = 0.0034). However, this cell-based assay could not exclude the contribution of cellular adaptor proteins.

To directly test JCAD-F-actin binding, we reconstituted actin filaments *in vitro* using purified human platelet actin. Staining of these reconstituted actin filaments with the mEmerald-JCAD protein revealed striking association (Figure 4G). Analysis of signal correlation revealed mEmerald-JCAD signal to be significantly more correlated with the actin fibers than the mEmerald control (Figure 4H; p < 0.0001). Together these data define JCAD as a novel molecular link between the TJ plaque and the actin cytoskeleton.

### JCAD forms dynamic condensates that associate with F-actin and co-partition with ZO-1

Recent work has established that TJ assembly involves liquid-liquid phase separation of ZO proteins (Pombo-García et al., 2024). Actin filaments contribute to this process by cross-linking

the ZO condensate network, and disruption of actin polymerization fragments the junctional ZO belt into discrete liquid-like droplets that reassemble upon actin repolymerization (Beutel et al., 2019; Schwayer et al., 2019). Since JCAD localizes along actin fibers in a punctate pattern, binds F-actin directly, and strongly co-localizes with ZO-1 at junctions, we asked whether JCAD participates in actin-associated TJ condensates.

At both endogenous and overexpressed levels, JCAD displayed a punctate distribution along actin fibers reminiscent of the droplet-like structures described for ZO proteins (Figures 4C & 4E) (Beutel et al., 2019; Schwayer et al., 2019; Pombo-García et al., 2024). To examine the dynamics of these structures, we performed live-cell imaging of hMVECs co-expressing JCAD-mEmerald and Lifeact-mScarlet. JCAD associated with actin filaments in a discontinuous pattern, accumulating in discrete puncta that moved along filaments and frequently underwent both fusion and fission events (Figure 5A).

**Figure 5:**
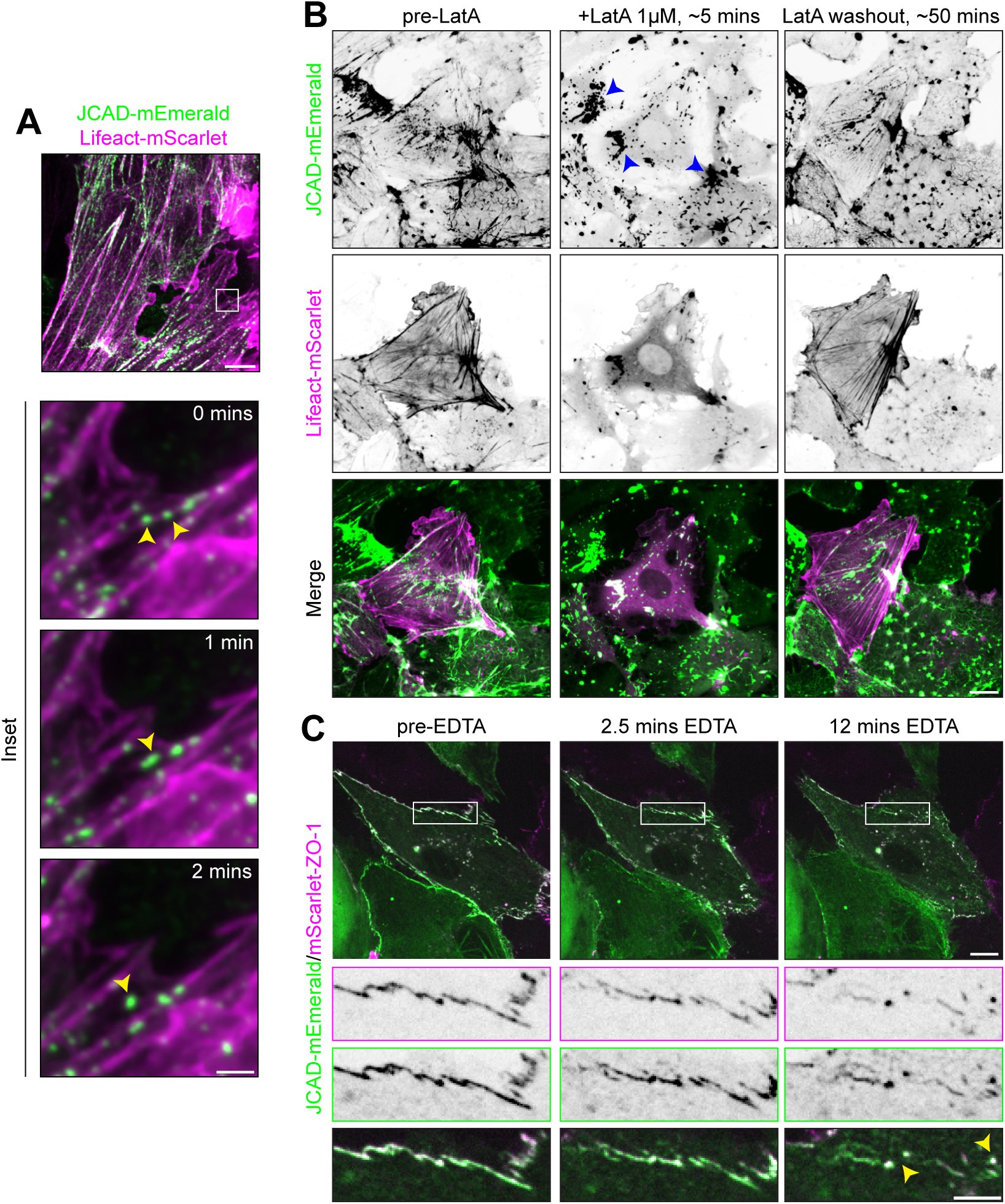
JCAD droplets associate with F-actin and co-partition with ZO-1. **(A)** Live-cell imaging of hMVECs co-expressing JCAD-mEmerald and Lifeact-mScarlet reveals fusion and fission of JCAD-mEmerald puncta. Scale bar, 5 µm. White box, location of inset (scale bar, 1 µm). Yellow arrowheads, puncta fusion event. **(B)** Live-cell imaging of hMVECs co-expressing JCAD-mEmerald and Lifeact-mScarlet before, 5 minutes after LatrunculinA treatment (1 µM), and 50 minutes after LatrunculinA washout. Blue arrowheads, examples of large JCAD aggregates after actin depolymerization. Scale bar, 15 µm. **(C)** Live-cell imaging of hMVECs co-expressing JCAD-mEmerald and mScarlet-ZO-1 treated with 2.5 mM EDTA reveals co-partitioning of JCAD and ZO-1 droplets during junctional disassembly. Scale bar, 10 µm. White box, location of inset (scale bar, 5 µm). Yellow arrowheads, JCAD-ZO-1 co-partitioned droplets.

Because actin filaments appeared to organize JCAD puncta, we next tested whether actin polymerization influences their distribution. Treatment with Latrunculin A, which sequesters monomeric actin and disrupts filament assembly, caused rapid redistribution of JCAD-mEmerald into large aggregated condensates reminiscent of the ZO-1 droplet formation (Beutel et al., 2019) observed upon actin disruption (Figure 5B, +LatA). Importantly, following Latrunculin A washout and repolymerization of the actin network, JCAD progressively localized from these aggregates back onto actin filaments (Figure 5B, LatA washout; Video 3). The reversibility of this process upon actin repolymerization confirms that JCAD puncta are dynamic condensates rather than irreversible aggregates.

Given the strong spatial correlation between JCAD and TJ components (Figure 3), we next examined whether JCAD associates with ZO-1 condensates during junction disassembly. Building on our earlier EDTA chelation experiments (Figure 3E-G), we examined JCAD-mEmerald and ZO-1-mScarlet localization at later timepoints when TJs were beginning to disassemble and condense into discrete puncta. Live-cell imaging revealed that upon junctional disassembly, JCAD and ZO-1 coalesced into shared punctate structures (Figure 5C). Altogether, we observe that JCAD forms dynamic punctate assemblies that associate with actin filaments, respond reversibly to actin polymerization state, and co-partition with ZO-1 during junctional remodeling.

### JCAD associates with tight junctions and actin through previously uncharacterized functional domains

JCAD harbors few reported functional motifs beyond tandem PPxY motifs (Lin et al., 2019), a pLxIS motif (Landau et al., 2024), and a short region of limited homology to ROCK1/2 and cingulin (Figure 6A, top) (Akashi et al., 2011). To identify additional functional domains underlying JCAD regulation of endothelial adhesion and barrier function, we undertook a combined structural prediction and evolutionary conservation analysis.

**Figure 6:**
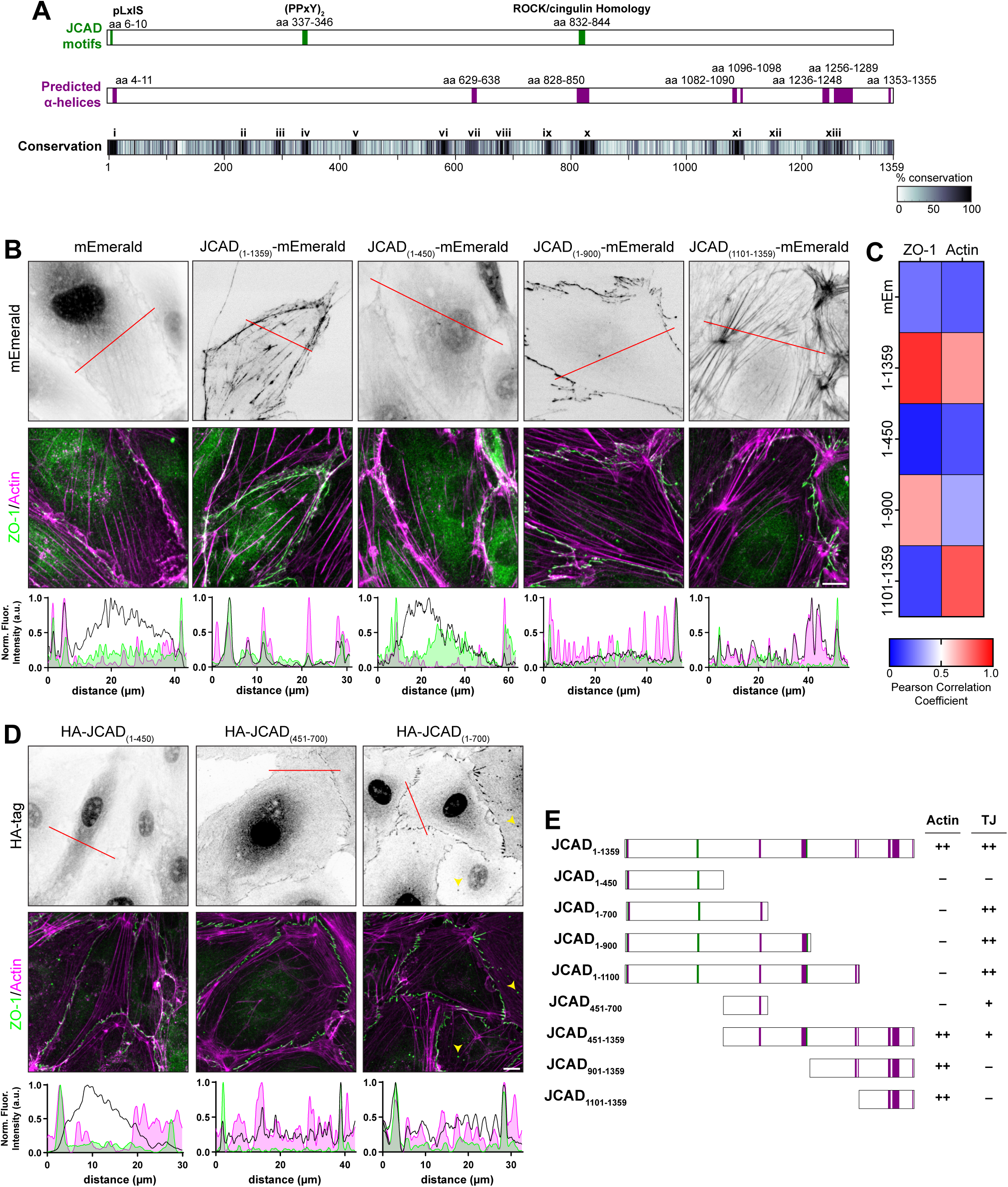
Novel JCAD functional domains revealed by phylogenetic analysis and truncation analysis. **(A)** Schematic of JCAD known motifs (top), AlphaFold3 predicted α-helices (middle), and evolutionary conservation (bottom). All unlabeled regions in the middle panel are predicted to be unstructured. Highly conserved regions are labeled i through xiii. **(B)** Immunostaining for ZO-1 and actin in hMVECs expressing mEmerald or mEmerald-tagged JCAD truncations in a JCAD KD background. Red lines, location of line-scan intensity profile. Scale bar, 10 µm. **(C)** Quantification of Pearson correlation coefficient between ZO-1/F-actin and mEmerald-JCAD truncations. Average of three line-scans per JCAD truncation construct is displayed. **(D)** Immunostaining for ZO-1 and phalloidin labeling of F-actin in hMVECs expressing HA-tagged JCAD truncations. Red lines, location of displayed line-scan intensity profile. Yellow arrowheads, examples of JCAD and ZO-1 enriched non-junctional aggregates. Scale bar, 10 µm. **(E)** Summary of JCAD truncations and identified targeting domains.

AlphaFold3 structural modeling predicted JCAD to be predominantly disordered, with several discrete α-helical segments within extended unstructured regions (Figure 6A, middle) (Abramson et al., 2024). Because no clear functional domains were apparent, we hypothesized that evolutionarily conserved regions would identify functionally important interfaces. We therefore aligned all JCAD orthologs spanning vertebrate evolution, from *Petromyzon marinus* (sea lamprey) to *Homo sapiens*, and quantified residue conservation across aligned sequences. This analysis revealed multiple highly conserved regions interspersed with more variable segments. As a positive control, the tandem PPxY domain (conserved region *iv*; Figure 6A, bottom) exhibited 99.2% median conservation, consistent with its proposed signaling function (Lin et al., 2019). Together, these analyses identified candidate functional interfaces within JCAD.

### JCAD truncation mutants reveal distinct domains driving subcellular targeting

To determine whether the structured or conserved regions regulate JCAD subcellular localization, we generated rationally designed truncation mutants with breakpoints at amino acids 450, 900, and 1100, chosen to preserve predicted conserved and structured regions. We generated lentiviral vectors enabling simultaneous shRNA-mediated knockdown of endogenous JCAD and rescue with mEmerald-tagged truncations. Localization of full-length and truncated JCAD proteins in hMVECs was assessed relative to actin and ZO-1 by immunostaining and Pearson correlation analysis of intensity line scans.

Full length JCAD_(1-1359)_-mEmerald localized to both actin fibers and ZO-1 at cell-cell interfaces, as demonstrated by representative intensity line scans and high mean Pearson correlation coefficients, whereas mEmerald alone and JCAD_(1-450)_-mEmerald showed no specific localization (Figure 6B-C). In contrast, JCAD_(1-900)_-mEmerald and JCAD_(1-1100)_-mEmerald strongly co-localized with ZO-1, identifying a TJ-targeting region within the first 900 amino acids (Figure 6B-C; Figure S6A). Conversely, C-terminal fragments JCAD_(451-1359)_-mEmerald, JCAD_(901-1359)_-mEmerald, and JCAD_(1101-1359)_-mEmerald localized to actin, identifying an actin-associating region within amino acids 1101-1359 encompassing conserved regions xii-xiii and three predicted α-helices (Figure 6A).

To further refine the TJ-targeting domain, we expressed JCAD fragments spanning amino acids 1-450, 450-700, and 1-700. Consistent with earlier results, HA-JCAD_(1-450)_ did not localize to junctions, whereas HA-JCAD_(1-700)_-mEmerald localized similarly to JCAD_(1-900)_-mEmerald (Figure 6D), indicating that sequences between amino acids 700-900 – including conserved regions ix-x and a predicted α-helix (amino acids 828-850) – are dispensable for junctional targeting. Interestingly, HA-JCAD_(451-700)_ showed detectable accumulation at cell-cell interfaces, but this localization was weaker than HA-JCAD_(1-700)_ and lacked the punctate pattern observed for ZO-1.

Notably, HA-JCAD_(1-700)_ also localized to non-junctional ZO-1-positive puncta (Figure 6D, arrowheads). High-resolution imaging revealed hollow, sphere-like JCAD condensates coated non-uniformly by ZO-1 (Figure S6B), consistent with partial co-partitioning between JCAD and ZO-1 assemblies. Together, these data suggest that amino acids 1-450 promote efficient condensate partitioning, whereas amino acids 450-700 are sufficient for interaction with junctional components.

Collectively, these data support a model in which JCAD is organized into at least two functional modules: an N-terminal region mediating TJ localization and junctional assembly association, and a C-terminal region promoting actin association (Figure 6E). This organization suggests a mechanism by which JCAD may couple the junctional plaque to the actin cytoskeleton to support cell-cell adhesion integrity.

### JCAD couples the tight junction to RhoA signaling

We next asked how JCAD facilitates barrier regulation during inflammatory challenge. JCAD is among the most highly and rapidly phosphorylated proteins after endothelial cells are treated with thrombin, a pro-inflammatory RhoA agonist that disrupts vascular barrier integrity, suggesting that JCAD may regulate barrier function through RhoA or its downstream effectors (Van Den Biggelaar et al., 2014). The presence of a ROCK homology region within JCAD further supports this link (Figure 6A), raising the possibility that JCAD functions as a local scaffold coupling TJs to actomyosin contractility.

To assess the role of JCAD in regulating RhoA-mediated inflammatory responses, we quantified thrombin-induced barrier disruption by measuring exposed substrate area following acute stimulation. Confluent monolayers of Control and JCAD KD cells were plated on biotinylated fibronectin-coated coverslips and treated with thrombin (2 U/mL, 2 min) in the presence of fluorescently labeled streptavidin. JCAD KD cells exhibited a significantly greater increase in exposed area, characterized by the formation of large intercellular gaps (Figure 7A-B). These findings are consistent with our *in vivo* results and suggest that JCAD limits inflammatory barrier responses downstream of RhoA.

**Figure 7:**
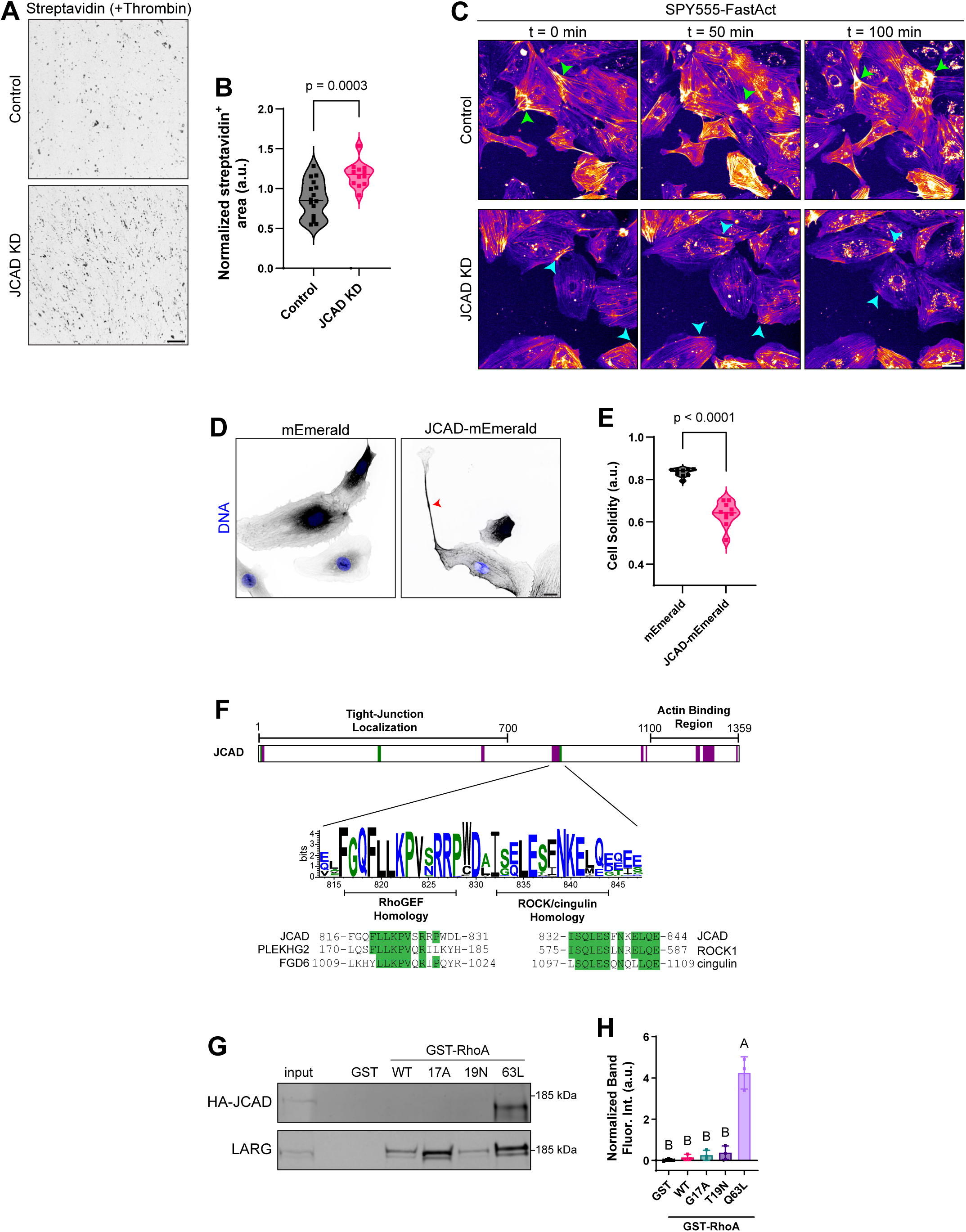
JCAD couples the tight junction to RhoA signaling. **(A)** Representative images of exposed substrate, labeled by Alexa Fluor conjugated streptavidin, after Control and JCAD KD cells were treated with thrombin (2 U/mL, 2 min). Scale bar, 200 µm. **(B)** Quantification of normalized streptavidin positive area (N = 3 independent experiments; n = 4-8 fields of view per N; Welch’s *t*-test). **(C)** Representative live-cell images demonstrating F-actin organization in Control and JCAD KD cells. Actin visualized using SPY555-FastAct probe. Green arrowheads, actin clusters exhibiting local contractility. Cyan arrowheads, nascent, unproductive actin clusters. Scale bar, 40 µm. **(D)** Representative images of hMVECs expressing mEmerald and mEmerald-JCAD. Scale bar, 15 µm. **(E)** Quantification of cellular solidity for mEmerald and mEmerald-JCAD overexpressing hMVECs (N = 3 independent experiments; n = 3-4 fields per N; Welch’s *t*-test). **(F)** Schematic demonstrating JCAD ROCK/cingulin homology domain and newly identified homology to RhoGEF-domain containing proteins. **(G)** Western blot for HA-tag and LARG (positive control) in RhoA mutant GST affinity precipitation assays. **(H)** Quantification of HA-JCAD western blot band intensity. Groups not sharing a letter are significantly different (N = 3 independent experiments; one-way ANOVA with Tukey’s post hoc test).

We next investigated whether JCAD regulates the actin dynamics associated with RhoA signaling. Live-cell imaging of F-actin in JCAD KD cells revealed a distinct actin architecture relative to Control cells. In Control cells, cortical actin puncta – previously shown to regulate endothelial mechanics during angiogenesis (Mayo et al., 2026) – were readily observed and correlated with local actin network contraction (Figure 7C, green arrowheads). In JCAD KD cells, these mesoscale structures were markedly reduced, and when present, were less persistent and failed to drive coordinated inward contractility toward the clusters (Figure 7C, cyan arrowhead; Video 4). Consistent with JCAD regulating RhoA-mediated actin dynamics, live-cell phase-contrast imaging revealed that JCAD KD hMVECs exhibited significantly impaired migration relative to Controls, as measured by both migration velocity and confinement ratio (Figure S7).

Conversely, JCAD-overexpressing cells revealed an opposing phenotype: an aberrant spindle-like morphology with processes extending from the cell body (Figure 7D, arrowhead). Cellular solidity (cell footprint area/convex hull area) of JCAD-overexpressing cells was significantly reduced compared to mEmerald-expressing controls (Figure 7E). Together, these data are consistent with JCAD influencing actin dynamics and cell shape directed by RhoA signaling and demonstrate that precise JCAD levels are required to maintain endothelial mechanical homeostasis.

Re-examination of conserved JCAD homology domains (Figure 6A) revealed that the 13 amino acid ROCK/cingulin homology domain (amino acids 832-844) exhibits a median residue-wise conservation of 84%, while an extended flanking region (amino acids 816-841) showed markedly higher conservation at 99%, suggesting that this broader region may encode a functionally important interface. BLAST analysis of amino acids 816-832 revealed homology to the RhoGEF domains of multiple Rho guanine exchange factors, including PLEKHG1, PLEKHG2, FGD6, and FARP2 (Figure 7F), raising the possibility that JCAD functions as a novel RhoA binding partner at TJs.

To test this, we purified recombinant GST-tagged RhoA bearing point mutations that stabilize the active GTP-bound (Q63L), inactive GDP-bound (T19N), or nucleotide-free (G17A) conformations. HA-tagged JCAD was expressed in hMVECs, and GST affinity precipitation was performed followed by western blot analysis of precipitated proteins. Whereas leukemia-associated RhoGEF (LARG), a known RhoA-binding positive control, exhibited binding to all RhoA conformations, JCAD demonstrated robust and selective binding to active RhoA(Q63L) only, with no appreciable binding to the other conformations (Figure 7G-H).

Collectively, these data demonstrate that JCAD loss sensitizes endothelial cells to RhoA-mediated barrier disruption and that JCAD selectively associates with GTP-RhoA. The heightened intercellular gap formation observed in JCAD KD cells following thrombin stimulation mirrors the inflammation-dependent vascular hyperpermeability of *Jcad^-/-^* mice, consistent with a shared mechanism regulating endothelial adhesion mechanics downstream of RhoA. Altogether, this suggests that JCAD serves as a scaffold of active RhoA, directing its activity toward TJ-associated actin to maintain the endothelial barrier.

## Discussion

Here we identify JCAD as a modular scaffold that organizes endothelial TJ architecture by coupling ZO-1-associated condensates to the actin cytoskeleton and RhoA signaling. Although *JCAD* has been genetically linked to vascular disease, a molecular function at endothelial cell-cell adhesions has remained unclear. Our findings define JCAD as a TJ-associated scaffold required to maintain junctional integrity during inflammatory stress, providing a mechanistic link between condensate-based TJ organization and actin regulation in the endothelium.

A key conclusion of this study is that JCAD is a component of the TJ plaque, and that its loss causes junction-type-specific effects: JCAD KD selectively disrupts TJ, but not AJ, continuity, establishing that JCAD regulation of paracellular permeability is mediated through the TJ. While prior studies reported JCAD localization to be VE-cadherin dependent (Akashi et al., 2011), we demonstrate that JCAD localizes to ZO-1-positive junctions independently of VE-cadherin and remains junctional upon acute AJ disruption. Together with a previously published ZO-1 proximity labeling experiment identifying JCAD near the phase separation-promoting ZO-1 C-terminus (Beutel et al., 2019; Van Itallie et al., 2013), these data support a revised model in which JCAD is integrated into the TJ plaque, and that junction-type-specific scaffolding underlies its role in vascular barrier maintenance.

That JCAD was nonetheless robustly labeled in our VE-cadherin BioID screen may reflect the presence of a stable, immobile pool of VE-cadherin anchored to the cortical actomyosin cable in close proximity to the actin-anchored TJ. Consistent with this, JCAD was also identified in a VE-cadherin BioID screen in lymphatic endothelial cells treated with adrenomedullin (Serafin et al., 2024), which stabilizes lymphatic barrier and cell-cell junctions through Rac1-mediated cortical actin polymerization (Koyama et al., 2013; Dunworth et al., 2008). While the characteristics described here implicate JCAD as a TJ protein, whether JCAD plays a cross-functional role spanning both junction types remains an open possibility.

Our data support a bipartite scaffold model in which JCAD couples TJ condensates to the actin cytoskeleton through distinct domains. It is notable that as expression levels increase, JCAD progressively localizes first to cell-cell contacts and then to actin filaments. Redistribution of a TJ protein to actin fibers across the cell body specifically in the overexpression context has likewise been described for junction-associated coiled-coil protein (JACOP), also known as cingulin-like 1 (CGNL1) (Ohnishi et al., 2004). Consistent with JCAD facilitating local control of RhoA-mediated actin network organization, we find that JCAD KD cells display a marked reduction in cortical actin puncta and, when these structures do form, they are shorter-lived and fail to drive coordinated contractility. Together, these findings position JCAD as an organizational node that integrates TJ condensates and the actin cytoskeleton to preserve endothelial barrier integrity under stress.

Beyond its structural role, we find that JCAD contains a short region with homology to proteins containing RhoGEF domains, and that JCAD biochemically associates with RhoA in the GTP-bound conformation, but not in the nucleotide-free or GDP-bound conformation. RhoA is known to drive cellular contractility and barrier breakdown in response to inflammatory stimuli, yet is equally critical for local repair of junctional breaks (Stephenson et al., 2019), implying that spatially organized scaffolds must direct RhoA activity in a context-dependent manner. JCAD is well positioned to mediate this spatial regulation; however, whether it does so by modulating RhoA guanine nucleotide exchange, GTPase activity, or by acting as a novel downstream effector remains to be determined. Nonetheless, JCAD-RhoA-GTP binding specificity – combined with JCAD’s structural capacity to act as a TJ-actin tether – raises the possibility that JCAD spatially directs RhoA activity at the TJ to locally coordinate actomyosin dynamics during barrier maintenance and remodeling. Intriguingly, the primary cellular function previously attributed to JCAD is regulation of Hippo signaling; however, Hippo pathway activation by RhoA agonists, such as TRAP6, was shown to require JCAD (Jones et al., 2018). Together with our findings, these data indicate that JCAD-dependent effects on Hippo signaling may be secondary to its regulation of RhoA and that JCAD may integrate cytoskeletal and transcriptional responses at the TJ.

Collectively, this model helps explain the stress-dependent requirement for JCAD in vascular barrier maintenance. JCAD*-*deficient mice maintain normal baseline barrier integrity but exhibit increased vascular leakage under endotoxemia-induced inflammatory conditions. Stress-dependent barrier effects are common in mice carrying genetic alleles that affect endothelial function, for example in PECAM1-null (Carrithers et al., 2005) and VE-cadherin mutant mice (Grimsley-Myers et al., 2020). During endotoxemia, endothelial junctions are subjected to increased mechanical and inflammatory stress, necessitating rapid and coordinated cytoskeletal remodeling to maintain barrier function. The rapid phosphorylation of JCAD in response to thrombin treatment is a plausible mechanism by which JCAD may be regulated to modulate vascular barrier. By coupling TJ structure to RhoA-mediated actomyosin dynamics, JCAD may enable this adaptive response; upon its loss or impairment, this coordination fails, resulting in junctional discontinuities and barrier leak.

Finally, genetic associations linking JCAD to coronary artery disease (Peden et al., 2011; Erdmann et al., 2011) and late-onset Alzheimer’s disease (Murdock et al., 2013) suggest that its junctional function may have broad relevance to vascular pathology, as both conditions are strongly associated with chronic vascular barrier dysfunction (Hahn and Schwartz, 2009; Zlokovic, 2011). Our results provide a cellular framework by which impaired JCAD-dependent junctional coupling could increase susceptibility to chronic endothelial dysfunction. Beyond pathologies clearly linked to vascular dysfunction, JCAD has also been described as a candidate tumor suppressor in serous borderline tumors of the ovary (Boyd et al., 2013) and high JCAD expression is associated with poor outcomes in cervical and stomach cancers (Uhlen et al., 2017; Yuan et al., 2025), suggesting that these novel functions for JCAD may be applicable for other cells beyond the endothelium.

In summary, we identify JCAD as a modular scaffold that integrates ZO-1-associated TJ condensates with actin dynamics and RhoA signaling to maintain endothelial barrier integrity during inflammatory stress. These findings establish a framework for understanding how phase-separated junctional assemblies are coupled to mechanical signaling in the endothelium and suggest that disruption of this integration may represent a general mechanism of vascular dysfunction.

## Materials and Methods

### Antibodies and reagents

Antibody and reagent information are provided in Table S1.

### Cell Culture

All cells were maintained in a humidified incubator at 37 °C with 5% CO_2_ and were tested for mycoplasma (Applied Biological Materials). The following cells were maintained in EGM2-MV medium (Lonza; CC-3202) and were used for experiments between passages two and ten: (1) human dermal microvascular endothelial cells (hMVECs; Lonza; CC-2813; lot 7F3585), (2) human coronary artery endothelial cells (hCAECs; Lonza; CC-2585; lot 20TL065655), (3) human brain microvascular endothelial cells (Angio-Proteomie; cAP-0002), and (4) human dermal lymphatic endothelial (hDLEC; Lonza; CC-2543). Human umbilical vein endothelial cells (hUVECs; Lonza; C2519A) were maintained in EGM2 (Lonza; CC-3162). Fluid shear stress was applied to cells where indicated using an orbital shaker following published protocol (Singh et al., 2026).

Caco-2 were a gift from the Tejal Desai Laboratory (UCSF) and were maintained in Dulbecco’s Modified Eagles Medium (DMEM) supplemented with 10% heat-inactivated fetal bovine serum (FBS; Peak Serum or SeraPrime), 1x MEM Non-Essential Amino Acids (Gibco; 11140050), 100 U/mL penicillin, and 100 µg/mL streptomycin (Sigma-Aldrich; P4333). U-2 OS were a gift from the Christopher Chen Laboratory (Boston University) and were maintained in DMEM supplemented with 10% heat inactivated FBS, 1 mM sodium pyruvate (Gibco; 11360070), 100 U/mL penicillin, and 100 µg/mL streptomycin (Sigma-Aldrich; P4333). HEK-293T (Clonetech) were maintained in DMEM supplemented with 10% heat inactivated FBS, 1x MEM Non-Essential Amino Acids (Gibco; 11140050), 10 mM HEPES (Gibco; 15630080), 1 mM sodium pyruvate (Gibco; 11360070), 100 U/mL penicillin, and 100 µg/mL streptomycin (Sigma-Aldrich; P4333).

#### Endothelial barrier measurement

The assay was conducted as previously described (Rathod et al., 2024; Aw et al., 2025) with minor modifications. To prepare biotinylated fibronectin, human fibronectin (Corning; 354008) was resuspended in ice-cold ddH_2_O to 1 mg/mL. EZ-Link Sulfo-NHS-LC-LC-Biotin (Thermo Fisher Scientific; 21338) was equilibrated to room temperature and resuspended to 10 mM in ddH_2_O and immediately added to fibronectin diluted to 0.1 mg/mL in PBS, yielding a final biotin concentration of 0.5 mM. The reaction was incubated at room temperature for 30 min. The resulting product was aliquoted and stored at -70 °C.

To coat coverslips, biotinylated fibronectin was thawed on ice and diluted 1:1 with PBS to 50 µg/mL. 25mm glass coverslips were plasma-cleaned for 30 s at 300 mTorr and incubated with the fibronectin solution for 60 min at 37 °C in a humidified incubator. Coverslips were washed three times with sterile PBS and UV sterilized for 15 min.

Cells were seeded onto biotinylated fibronectin-coated coverslips and grown to confluency. Media was replaced, and fluid shear stress was applied using an orbital shaker as previously described (Singh et al., 2026). All subsequent reagents were equilibrated to 37 °C prior to use. After 24 h of culture under shear, media was aspirated and coverslips were incubated with 25 µg/mL AlexaFluor-conjugated streptavidin (Invitrogen; S11226) in PBS containing calcium and magnesium (PBS^++^; Gibco; 14040182) for 2 min. Coverslips were washed twice with PBS^++^ and fixed in 4% (w/v) paraformaldehyde in PBS^++^ for 10 min at 37 °C.

Fixative was aspirated and coverslips were washed three times with PBS before incubation in a quenching/blocking buffer (PBS containing 100 mM glycine and 2% [w/v] BSA) for 1 hr at room temperature. Coverslips were then incubated with FITC-conjugated mouse anti-VE cadherin for 1 hr at room temperature, washed three times with PBS, and mounted using ProLong Glass (Invitrogen). For quantification of streptavidin positive area, coverslips were imaged at 4x magnification.

#### Thrombin response assay

JCAD KD and Control cells were plated on coverslips coated with biotinylated fibronectin as described for endothelial barrier measurements. Thrombin (2 U/ml; Sigma Aldrich; 604980) was added to the PBS/streptavidin solution and after two minutes of treatment, coverslips were processed as described for barrier measurements.

#### Live-cell imaging

For live-cell imaging experiments, cells were plated on glass-bottom dishes coated with 50 µg/mL rat tail collagen I. For imaging of actin dynamics, cells were incubated with SPY555-FastAct (1:2000) for ∼2 hours prior to imaging. Latrunculin A (1 µM) or EDTA (2.5 mM) was added directly to the culture media during acquisition. For latrunculin A washout experiments, media was aspirated and cells were quickly washed once with fresh media before continuing imaging. For doxycycline-treated conditions, doxycycline (2 µg/mL) was added to media immediately prior to imaging.

### Microscopy

Images in this study were collected on one of three spinning disc confocal microscopes.

The first scope was a Yokogawa CSU10 spinning disk confocal on a Nikon TE2000 microscope with a CoolSNAP HQ2 camera (Photometrics). 405 nm, 488 nm, 561 nm, and 640 nm solid-state lasers were used at powers between 10 and 100%.

The second scope was a Yokogawa CSU-X1 spinning disk confocal custom-modified by Spectral Applied Research on a Nikon Ti-E microscope with an Evolve EMCCD camera (Photometrics) (Stehbens et al., 2012). 100 mW diode-pumped solid-state 488 nm (Coherent Sapphire) and 561 nm (Cobolt Jive) lasers were used at powers between 10 and 100%. The imaging system was enclosed within an imaging chamber (In Vivo Scientific) at 37 °C with humidified air at 5% CO_2_ to facilitate live-cell imaging.

The third scope was a Yokogawa CSU-W1/SoRa spinning disc confocal system and an ORCA Fusion BT camera (Hamamatsu). A Nikon 60 × NA 1.49 oil immersion objective lens was used with 100 mW lasers (405 nm, 488 nm, 561 nm, 647 nm). The scope was equipped with a stage-top incubator (Okolab) at 37 °C with humidified air at 5% CO_2_ to facilitate live-cell imaging. All scopes were controlled using NIS Elements software (Nikon).

### Lentivirus production and transduction

HEK 293T were plated at a density of 7×10^6^ cells per 100 mm dish or 8×10^5^ cells per well of a 6 well plate. The following day, cells were transfected with pMD2.g, psPAX2, and the transfer plasmid in a molar ratio of 1:2:3, respectively using calcium phosphate transfection. After ∼6 hours, the media was changed. Two days later, the media was harvested, either filtered through a 0.2 µm syringe filter or centrifuged at 800 x g for 10 minutes, and concentrated overnight at 4°C using a 3× PEG solution (40% [w/v] PEG-8000 and 1.2 M NaCl). Viral particles were pelleted by centrifugation (1600 × g for 1 hour at 4 °C) and resuspended in PBS.

Target cells were transduced with lentivirus immediately after passaging and were incubated with virus overnight (12-18 hours). The following day, the media was replaced. For constructs containing a puromycin resistance cassette, cells were selected for up to two days using 2 µg/mL puromycin (Sigma-Aldrich; P8833).

### Vertebrate Animal Experiments

All animal experiments were performed in accordance with protocols approved by the University of California, San Francisco Institutional Animal Care and Use Committee.

#### Generation of Jcad null allele

*Jcad null allele (Jcad^-^)* was generated using the CRISPR/Cas9-based method called i-GONAD (Gurumurthy et al., 2019; Ohtsuka et al., 2018) by sequential deletion of two exons. CRISPR guide RNAs were designed using the Integrated DNA Technologies gRNA design tool. To delete exon 2 of *Jcad (Jcad^Δexon2^)*, C57BL/6J wildtype males were crossed with CD1 wildtype females. At embryonic day 0.7 (E0.7), oviducts of pregnant females were injected with 0.8 μL of a mixture containing 0.03 nmol of each CRISPR RNA (crRNA1: 5’-AGTCCCTTACCTGTACCAAG-3’ and crRNA2: 5’-AAGCACTTCTAGAACTCCTG-3’), 0.06 nmol trans-activating crRNA (tracrRNA), and 2 μg Cas9 protein, followed by electroporation. Germline transmission of an edited allele was confirmed by sequencing the progeny of a candidate founder. The allele was maintained on a mixed CD1/C57BL/6J background and backcrossed three generations to C57BL/6J.

To delete exon 3 of *Jcad (Jcad^Δexon3^),* heterozygous intercross of N3 *Jcad^+/Δexon2^* mice yielded homozygous males (*Jcad^Δexon2/Δexon2^*) which were crossed to wildtype CD1 females. The oviducts of pregnant females were injected as previously described (crRNA3: 5’-GGGTATCCTCACCTCAACGC-3’ and crRNA4: 5’-GGTTACCACCTTTGAAACAT-3’). Again, germline transmission of an edited allele was confirmed by sequencing the progeny of a candidate founder.

Upon first backcross of *Jcad^Δexon3^* founder to C57BL/6J wildtype mice, *Jcad^Δexon2^* and *Jcad^Δexon3^* were co-inherited in the progeny confirming that these deletions were made on the same chromosome. The compound deletion allele (*Jcad^-^*) was maintained on a mixed CD1/C57BL/6J background and backcrossed three generations to C57BL/6J before experiments were conducted.

#### Allele validation and genotyping

Tail biopsies were lysed in 50 mM NaOH by heating to 95 °C for 45 minutes. *Jcad^Δexon2^* deletion was validated by PCR amplification using the following primers: Jcad-ex2-WT_fwd, Jcad-ex2-mut_fwd, and Jcad-ex2_rev. Wildtype allele yields an amplicon of length ∼750 bp and *Jcad^Δexon2^* yields ∼450 bp. *Jcad*^-^ allele transmission was validated by PCR amplification using the following primers: Jcad-ex3_fwd, Jcad-ex3-mut_rev, and Jcad-ex3-WT_rev. Wildtype allele yields an amplicon of length ∼850 bp and *Jcad^Δexon3^* yields ∼1000 bp. PCR was conducted using Q5 High-Fidelity 2X Master Mix (New England Biolabs; M0492) with annealing temperature of 60 °C and 60 seconds extension. Primer sequences are listed in Table S2.

#### Mouse vascular permeability assay

Mice were injected intraperitoneally with either 10 mg/kg body weight lipopolysaccharides from Escherichia coli O111:B4 (1 mg/mL; Sigma-Aldrich; L4391) or sterile saline. After three hours, mice were anesthetized using ketamine (100 mg/kg body weight; Dechra) and xylazine (10 mg/kg body weight; Dechra) and were injected with 100 µl Evans blue (1% [w/v] in sterile saline) retroorbitally. After 15 min of Evans blue circulation, mice were perfused through the left ventricle with 20 mL sterile PBS. Lung and brain tissue were harvested and Evans blue was extracted by incubating tissue with formamide overnight at 50°C. Extracted Evans blue was quantified by spectrometric analysis of absorbance at 620 nm (A620) and was normalized to the dry weight of the tissue from which it was extracted.

### Molecular biology

Plasmids used in this study are listed in Table S1 and were generated herein unless otherwise stated. All primers used are listed in Table S3. Constructs were sequence validated (Plasmidsaurus or Quintara Biosciences).

To generate hMVEC cDNA, hMVECs were lysed in TRI Reagent (Sigma Aldrich; T9424), RNA was isolated using the Direct-zol RNA Miniprep Kit (Zymo Research; R2050), and reverse transcription was conducted using SuperScript III Reverse Transcriptase (Invitrogen; 18080044). Polymerase chain reaction was conducted using Q5 High-Fidelity 2X Master Mix (New England Biolabs; M0492). Gibson Assembly was conducted using NEBuilder HiFi DNA Assembly Master Mix (New England Biolabs; E2621). shRNA sequences were ligated into modified pLL3.7 vectors using T4 DNA Ligase (New England Biolabs; M0202L).

pMD2.G and psPAX2 were gifts from Didier Trono (Addgene plasmids # 12259 and 12260). pCW57-MCS1-2A-MCS2 was a gift from Adam Karpf (Addgene plasmid # 71782). pLentiCRISPRv2 Scramble and CDH5 sgRNA were described previously (Polacheck et al., 2017). pGEX-6P-2 was a gift from Diane Barber Laboratory.

### Immunostaining

#### Hydrogel coverslip preparation

Polyacrylamide gels (2.5 kPa stiffness) were prepared and coated with 50 µg/mL rat-tail collagen I as previously described (Mayo et al., 2026).

#### Coverslip immunostaining

Coverslips were fixed in 4% (w/v) paraformaldehyde (Electron Microscopy Sciences) prepared in PBS containing calcium and magnesium (PBS^++^; 0.9 mM CaCl₂, 0.49 mM MgCl₂), pre-warmed to 37 °C, for 10 min. Fixed coverslips were washed once with PBS and quenched with 100 mM glycine in PBS for 30 min at room temperature. Cells were permeabilized with 0.1% (v/v) Triton X-100 in PBS for 10 min at room temperature, washed once with PBS, and blocked with 2% (w/v) bovine serum albumin in PBS for at least 1 h.

Primary antibodies (Table S1) were diluted in blocking solution and incubated with coverslips for 1-2 h at room temperature. Following three 10 min washes with PBS, coverslips were incubated with secondary antibodies and/or fluorescent dyes prepared in blocking solution for 1 h at room temperature. Coverslips were washed three times with PBS (10 min each), mounted using ProLong Glass Antifade Mountant (Invitrogen), sealed with nail polish, and stored at 4 °C until imaging.

#### Mouse tissue

Mice were anesthetized by intraperitoneal injection of ketamine (100 mg/kg body weight; Dechra) and xylazine (10 mg/kg body weight; Dechra) and were euthanized by transcardial perfusion of PBS. Retinas were dissected according to established protocol (Tual-Chalot et al., 2013). After dissection, retinas were fixed in ice cold 4% PFA (w/v) in PBS for 1 hour. After fixation, retinas were blocked and permeabilized in a PBS solution containing 0.3% (v/v) Triton X-100, 2% (w/v) bovine serum albumin, and 5% (v/v) normal goat serum overnight. Primary antibodies were diluted in blocking solution (Table S1) and incubated with retinas for 3 hours to overnight. Following three 10 min washes with PBS, retinas were incubated with secondary antibodies and/or dyes prepared in blocking solution for 1 h at room temperature. Retinas were again washed three times with PBS, and stored at 4 °C until imaging.

### Biochemistry

#### SDS-PAGE and Western Blot

Lysates from cultured cells were obtained by growing cells to 100% confluency. Cells were washed with ice cold PBS^++^, lysed in cold lysis buffer (pH 7.4, 25 mM Tris, 150 mM NaCl, 5 mM MgCl_2_, 1% v/v Triton X-100) with 2x Halt Protease and Phosphatase inhibitor (Thermo; 78442), and incubated on ice for 10 min. Lysates were centrifuged for 10 min at 4 °C at 21,380 × g. Supernatants were collected, snap frozen with liquid N2, and stored at -80 °C if not used immediately. Protein concentration was determined using a bicinchoninic acid reaction kit (Prometheus). Lysate was denatured using 1x NuPAGE LDS Sample Buffer (Invitrogen; NP0007) containing 5% β-mercaptoethanol for 10 min at 70 °C.

Lysates from mouse brains were obtained by dissecting tissue from mouse euthanized by CO_2_ inhalation. Brains were mechanically lysed in ice cold RIPA lysis buffer (pH 7.4, 50 mM Tris, 150 mM NaCl, 1% v/v Triton X-100, 0.1% w/v sodium dodecyl sulfate, 0.5% w/v sodium deoxycholate) containing 2x Halt Protease and Phosphatase inhibitor (Thermo; 78442). Lysates were centrifuged for 10 min at 4 °C at 21,380 × g and processed the same as cell culture lysates.

Denatured lysate of equalized protein content and PageRuler Plus prestained protein ladder (Thermo; 26619) were loaded into Invitrogen NuPAGE BisTris protein gels and run at 140-160V in 1x MOPS/SDS running buffer (20x MOPS/SDS buffer comprised of 1 M MOPS, 1 M Tris base, 69.3 mM sodium dodecyl sulfate, 20.5 mM EDTA free acid). Proteins were transferred from the gel onto nitrocellulose membrane in ice cold 1 × transfer buffer with 10% methanol (20x transfer buffer comprised of 500 mM bicine, 500 mM Bis-Tris, 20.5 mM EDTA free acid) using a Mini Trans-Blot Cell (Bio-Rad) at 4 °C. Membranes were blocked in 5% non-fat milk (Apex) or 5% bovine serum albumin in TBST (1x TBS containing 0.1% v/v Tween-20). Primary antibody was diluted in blocking solution and incubated for 1-2 h at room temperature with gentle rocking. Membranes were washed 3x in TBST for 10 min each with gentle rocking. Secondary antibody was diluted in blocking solution and incubated for 1 h at RT with gentle rocking. Membranes were washed 3x in TBST for 10 min each with gentle rocking. Membranes were imaged on an Odyssey CLx LI-COR Imaging System and quantified using Fiji/ImageJ.

For quantification of recombinant protein yield, samples of recombinant protein were subjected to SDS-PAGE with additional lanes containing known masses of bovine serum albumin. Polyacrylamide gels were washed with water three times to remove SDS and were stained with SimplyBlue SafeStain (Invitrogen; LC6060). Gel was imaged using an Odyssey CLx LI-COR Imaging System and band intensities were quantified using Fiji/ImageJ. Recombinant protein yield was quantified by comparing the band intensities to those of the bovine serum albumin calibration curve.

#### GST-RhoA Affinity Precipitation

hMVECs expressing HA-tagged JCAD (HA-JCAD) were grown to 100% confluency and lysates were collected as described with the addition of needle lysis prior to centrifugation (20 strokes through 25 G needle).

Sepharose beads coated in GST-RhoA proteins were thawed on ice and pre-equilibrated in lysis buffer. Each precipitation reaction contained approximately 20 µg GST-RhoA and approximately 600 µg cell lysate. Beads were incubated with lysates and rotated at 4 °C for 1 h. Beads were pelleted and washed with lysis buffer three times. After the third wash, all supernatant was aspirated and beads were resuspended in 2x NuPAGE LDS Sample Buffer (Invitrogen; NP0007) containing 5% β-mercaptoethanol and denatured for 10 minutes at 70 °C.

#### BioID Streptavidin Affinity Precipitation

hMVECs expressing a VE-cadherin-HA-tag-BirA* fusion protein (and wildtype control cells) were grown to 100% confluency in a 100 mm tissue culture dish and were treated with 50 µM biotin for 20 hours. After incubation, hMVEC monolayers were washed three times with ice cold PBS^++^ and were lysed in RIPA buffer (pH 7.4, 50 mM Tris, 150 mM NaCl, 1% v/v Triton-X, 0.1% w/v sodium dodecyl sulfate, 0.5% w/v sodium deoxycholate). Biotinylated proteins were extracted by incubating lysates with Dynabeads Streptavidin Magnetic Beads C1 (Invitrogen; 65001) for 1 hour at room temperature. Beads were washed three times with ice cold RIPA buffer and were resuspended in 2x NuPAGE LDS Sample Buffer (Invitrogen; NP0007) containing 5% β-mercaptoethanol and 50 µM biotin. Proteins were denatured for 10 minutes at 95 °C.

### Recombinant protein purification

#### GST-RhoA and mutants

pGEX plasmids were transformed into BL21(DE3) competent *E. coli* (Thermo Scientific; EC0114) and a single transformant of each line was stored as a glycerol stock (25% [v/v] glycerol) at -70°C. A starter culture of each glycerol stock was grown in 50 mL Luria broth medium (LB) containing 100 µg/mL carbenicillin overnight at 37 °C shaking at 225 RPM. The following day, the 50 mL culture was diluted into 450 mL LB containing 50 µg/mL carbenicillin and was incubated at 37 °C, shaking at 225 rpm, for 30 minutes. Protein expression was induced with 0.1 mM IPTG and cultures were incubated overnight at 20 °C with shaking.

Cells were harvested by centrifugation (4,000 × g, 15 min) and resuspended in lysis buffer (20 mM HEPES, pH 7.5; 150 mM NaCl; 5 mM MgCl₂; 1% [v/v] Triton X-100) supplemented with 1× Halt Protease Inhibitor Cocktail, EDTA-Free (Thermo Scientific; 78425) and 1 mM DTT. Cells were lysed by probe sonication (6 × 10 s pulses, 10 s rest, 8 W RMS, on ice) and insoluble material was removed by centrifugation (27,000 × g, 15 min, 4 °C).

Cleared lysates were incubated with 500 µL Glutathione Sepharose 4B (Cytivia; 17-0756-01) and rotated for 1 hour at 4 °C. Beads were washed three times with lysis buffer supplemented with 1 mM DTT, and then three times with 8 mL HEPES buffered saline (HBS; 20 mM HEPES, pH7.5; 150 mM NaCl) supplemented with 1 mM DTT and 5 mM MgCl_2_ (HBS^++^). Supernatant was removed to recover original bead volume and one half-volume of glycerol was added for cryoprotection. Beads were aliquoted, snap frozen in liquid nitrogen, and stored at -70 °C. Protein concentration was estimated by SDS-PAGE as described.

#### GST-mEmerald and GST-mEmerald-JCAD

Expression and lysis were performed as described for GST-RhoA proteins, but using Terrific Broth supplemented with 0.8% (v/v) glycerol. Protein expression was induced with 0.1 mM IPTG and 3% (v/v) ethanol while shaking 18 °C for ∼24 h.

Cleared lysates were incubated with 500 µL Glutathione Sepharose 4B (Cytivia; 17-0756-01) and rotated for 1 h at 4 °C. The beads were then pelleted and the mEmerald-JCAD supernatant was again incubated with 500 µL Glutathione Sepharose 4B to maximize yield. Protein-bound beads were washed three times with lysis buffer supplemented with 1 mM DTT, and then three times with HBS^++^. After the third wash, beads were pelleted, supernatant was removed, and beads were incubated with 10 mM reduced glutathione (Thermo; 78259) diluted in HBS^++^ for 15 minutes on ice. Supernatant containing the eluted protein was collected and this process was repeated to obtain a second elution fraction. Purified protein was stored at 4 °C for up to two weeks.

To validate purification, a small aliquot of each eluted sample was reduced, denatured, and subjected to SDS-PAGE. Gels were stained with Novex SimplyBlue SafeStain (Invitrogen; LC6060) and imaged using an Odyssey CLx LI-COR imaging system. Protein concentration was estimated by SDS-PAGE as described. Western blot analysis using a GFP antibody and a JCAD C-terminal antibody confirmed the presence of protein at expected molecular weights.

### *In vitro* actin binding assay

Actin polymerization was conducted using components of an Actin Binding Protein Spin-Down Assay Biochem Kit (Cytoskeleton, Inc.; BK013). General actin buffer (5 mM Tris-HCl pH 8.0 and 0.2 mM CaCl_2_) was resuspended in sterile water. The following reagents were resuspended according to the manufacturer’s instructions, snap frozen in liquid nitrogen, and stored at -70 °C: ATP (100 mM in 100 mM Tris, pH 7.5), human platelet actin (10 mg/mL in ice cold sterile water), BSA (3.4 mg/mL in sterile water), and actin polymerization buffer (500 mM KCl, 20 mM MgCl_2_, and 10 mM ATP in 100 mM Tris, pH 7.5).

To polymerize actin, an aliquot of 100 mM ATP was thawed, immediately placed on ice, and diluted to 10 mM ATP with ice cold sterile water, then further diluted to 0.2 mM ATP in general actin buffer. A 250 µg aliquot of actin was thawed quickly in a room temperature water bath and immediately moved to ice. Actin was diluted to 1 mg/mL using 225 µL general actin buffer containing 0.2 mM ATP and incubated on ice for 30 minutes. One aliquot of actin polymerization buffer was thawed and kept on ice. 25 µL of actin polymerization buffer was added to the actin tube, mixed by gentle pipetting, and incubated at room temperature for 1 hour yielding a 21 µM stock of F-actin.

Glass coverslips were plasma cleaned for 30 seconds at 300 mTorr and subsequently incubated with 0.01% (w/v) poly-L-lysine for 20 minutes at room temperature, followed by two washes with PBS. Coverslips were dried on Kimwipes, and a droplet of 21 µM F-actin was placed on a Parafilm sheet. Coverslips were inverted onto the droplet with the poly-L-lysine-coated surface contacting the F-actin and incubated at room temperature for 15 minutes to allow filament adsorption. Coverslips were fixed in 4% (w/v) paraformaldehyde for 3 minutes and washed twice with PBS containing 100 mM glycine. Fixed coverslips were incubated with recombinant GST-mEmerald or GST-mEmerald-JCAD at approximately equimolar concentrations (∼250 nM) for 15 minutes at room temperature, followed by three washes with PBS. F-actin was labeled with rhodamine-phalloidin (approximately 0.66 µM; Invitrogen; R415) for 5 minutes at room temperature. Coverslips were washed three additional times with PBS, mounted using ProLong Glass mounting medium, and imaged using a 100× objective.

### Phylogenetic conservation analysis

The amino acid sequence for one JCAD isoform from each ortholog was retrieved from NCBI. Protein sequences were analyzed in MATLAB (Mathworks) using Bioinformatics Toolbox functions. Sequences were aligned using progressive multiple sequence alignment based on pairwise distances calculated with the GONNET substitution matrix and guide trees generated by average-linkage clustering. *Homo sapiens* was selected as the anchor species, and alignment positions corresponding to gaps in the anchor sequence were removed to generate an anchor-referenced alignment. Residue-wise conservation was quantified by calculating the fraction of species sharing an identical amino acid at each aligned position, and conservation profiles were visualized as heatmaps. Sequence logo was generated using Web Logo 3.

### Image processing and analysis

Images were processed and analyzed using Fiji (Schindelin et al., 2012) and/or MATLAB (Mathworks) unless otherwise specified.

#### Streptavidin positive area

To minimize edge artifacts, images were cropped to a fixed central square region. Background fluorescence was subtracted using a rolling-ball algorithm (radius = 20 pixels). A single, fixed intensity threshold was applied across all images of each replicate, and binary masks were generated. The percentage of streptavidin-positive area was calculated as the area fraction of thresholded pixels relative to the total cropped image area.

#### Junctional skeletonization and discontinuity measurement

Images were cropped to a central region to minimize edge artifacts and junctions were identified by automated segmentation. The VE-cadherin channel was processed using sequential Gaussian blurring, top-hat filtering, and median filtering, followed by adaptive thresholding to generate a binary junction mask. Small particles were removed and remaining junctional structures were dilated to ensure continuity. The binary mask was then skeletonized, and individual junction segments were identified using the Analyze Skeleton (2D/3D) plugin (Doube et al., 2010) with shortest-branch pruning to remove spurious branches. Skeleton-derived ROIs were extracted and saved for reuse.

For each junction region of interest (ROI) isolated, area selections were converted to line ROIs and intensity line scans were extracted from the ZO-1 and VE-cadherin channels using fixed line widths (10 and 25 pixels, respectively). Junction length was recorded, and the coefficient of variation (CV) was calculated for each line scan. Per-junction line-scan data and summary statistics were exported and were further processed in MATLAB (Mathworks). For each junction, junction length and CV were used to compute a length-weighted variability metric (junction length × CV). Junctions were grouped by experimental condition, and outliers were excluded based on junctional CV using MATLAB’s built-in outlier detection. To account for unequal junction numbers per image, a per-image length-weighted mean CV was calculated by summing length-weighted CV values across all junctions within an image and normalizing by the total junction length for that image. These per-image values were used for statistical comparisons between conditions.

#### Junction protein IF correlation

Orthogonal intensity correlations at cell–cell junctions were quantified using a custom ImageJ/Fiji macro. Multichannel TIFF images were cropped to a central region, split into individual channels, and analyzed using pre-defined junction ROIs (see *Junctional Skeletonization*). For each junction, pixel coordinates along the ROI skeleton were used to generate orthogonal line profiles (±30 pixels) at each point along the junction. Pearson correlation coefficients were calculated for intensity values across orthogonal profiles between VE-cadherin-JCAD, and ZO-1-JCAD. Junctions with fewer than 400 orthogonal profiles were excluded. Mean correlation coefficients per junction were exported for downstream statistical analysis.

#### Junctional protein mean fluorescence intensity

Images were cropped to a central region to minimize edge artifacts. Cell-cell junctions were manually segmented, and mean fluorescence intensity was measured along each junction using a line ROI (width 50 pixels).

#### JCAD enrichment on junctional actin

Cell-cell junctions were manually cropped and binary masks were generated for both the JCAD and actin channels by applying intensity thresholds. To assess the spatial enrichment of JCAD on actin fibers, a randomized mask containing the same number of pixels as the JCAD mask was generated. The degree of colocalization was quantified by calculating the percentage of JCAD mask pixels overlapping with the actin mask, and this was compared to the percentage of overlap between the randomized mask and the actin mask. Enrichment was defined as the ratio of JCAD-actin overlap to random-actin overlap, such that a value greater than 1 indicates preferential association of JCAD with actin above chance.

#### GST-mEmerald/GST-mEmerald-JCAD and actin correlation

To minimize edge effects, the outer 15% of each image border was excluded by cropping to the central region. Pixel intensity values from the cropped regions were extracted for each channel (F-actin and mEmerald/mEmerald-JCAD), and Pearson correlation coefficients were computed on a per-image basis using all corresponding pixels.

#### JCAD truncation line-scan correlations

A straight-line ROI was manually drawn across the width of representative cells expressing the truncated proteins. Raw intensity profiles from junctional line scans were smoothed using a moving average filter (window = 10 pixels) prior to normalization and profiles were normalized to a 0-1 range by min-max scaling. Pearson correlation coefficient was calculated between the individual channels.

#### Cellular solidity

Images were thresholded based on the mEmerald or mEmerald-JCAD signal and boundaries between adjacent cells were manually annotated to create individual cell ROIs. The solidity (area divided by convex hull area) of each cell was measured and the average solidity for each field of view was recorded.

#### Cell migration metrics

Cell migration was assessed by manually tracking individual cell positions across time-lapse phase contrast images. For each cell, the tracked coordinates were used to calculate the total distance traveled, the net displacement, and mean velocity. The confinement ratio was calculated as the net displacement divided by the total distance traveled.

### Statistical Analyses

Statistical analyses were conducted in GraphPad Prism 11, with a significance threshold of α = 0.05 throughout. Data were tested for outliers using the ROUT method (Q = 1%), and identified outliers were excluded from analyses. Normality was assessed using the D’Agostino & Pearson test or the Shapiro-Wilk test, depending on sample size. For comparisons between two groups, normally distributed data were analyzed using Welch’s t-test, while non-normal data were analyzed using the Mann-Whitney U test. For comparisons of normalized data against a theoretical mean, a one-sample t-test was used. For comparisons between more than two groups, data were analyzed using one-way ANOVA with Tukey’s post hoc test, or two-way ANOVA with Holm-Šídák’s post hoc test when two independent variables were present.

## Acknowledgments

This work was supported by National Institutes of Health (NIH) grants R35GM150987 (MLK), R35DE031926 (JOB), F31HL172606 (LNM), R61CA278518 (TW), and S10OD028611 (TW; Nikon CSU-W1 SoRa). Other support was from the Leducq Foundation 21CVD03 (MLK), Tobacco-Related Disease Research Program Predoctoral Fellowship T33DT6442 (KAJ), and National Science Foundation Graduate Research Fellowship 2038436 (KAJ). The contents are solely the responsibility of the authors and do not necessarily represent the views of the NIH. Any opinions, findings, and conclusions or recommendations expressed in this material are those of the author(s) and do not necessarily reflect the views of the National Science Foundation. The monoclonal antibody developed by UC Davis/NIH NeuroMab Facility was obtained from the Developmental Studies Hybridoma Bank, created by the NICHD of the NIH and maintained at The University of Iowa, Department of Biology. Lung icon by Servier is licensed under CC-BY 3.0 Unported.

## Author Contributions

Conceptualization: MLK, KAJ

Data curation: KAJ

Formal analysis: KAJ

Funding acquisition: MLK, KAJ, JOB, LNM, TW

Investigation: KAJ, Y-GJ, F-SL, LNM

Methodology: KAJ, MLK, JOB

Project administration: MLK, KAJ, JOB, TW

Resources: KAJ, MLK, JOB, LNM

Supervision: MLK, JOB, TW

Validation: KAJ, MLK

Visualization: KAJ

Writing – original draft: KAJ

Writing – review & editing: KAJ, MLK, TW, JOB, Y-GJ, F-SL, LNM

## Data, code, and materials availability

Available from the corresponding authors upon reasonable request.

## List of Supplementary Materials

Figs. S1 to S7

Tables S1 to S3

Videos 1 to 4

---

**Video 1: Overexpressed mEmerald-JCAD dynamically localizes to a filamentous network at the trailing edge of migrating hMVECs.**

Live-cell imaging of hMVECs expressing mEmerald-JCAD under human cytomegalovirus promoter. Still frames annotated in Figure 4A. Scale bar, 20 µm.

**Video 2: Sequential localization of JCAD-mEmerald to cell-cell junctions and actin filaments.**

Live-cell imaging of hMVECs expressing JCAD-mEmerald under a doxycycline-inducible promoter following induction with doxycycline (2 µg/ml) at the start of imaging. At low expression levels, JCAD-mEmerald first localizes to cell-cell contacts and subsequently accumulates along actin filaments. Scale bar, 20 µm.

**Video 3: Actin depolymerization reveals JCAD phase separation and F-actin association.**

Live-cell imaging of hMVECs coexpressing JCAD-mEmerald and LifeAct-mScarlet before, during, and after Latrunculin A (LatA)-mediated actin depolymerization. Still frames annotated in Figure 5B. Scale bar, 15 µm.

**Video 4: JCAD depletion alters actin organization in endothelial cells.**

Live-cell imaging of control and JCAD KD hMVECs labeled with SPY555-FastAct to visualize filamentous actin organization. Still frames annotated in Figure 7C. Scale bar, 30 µm.

**Supplemental Figure S1:**
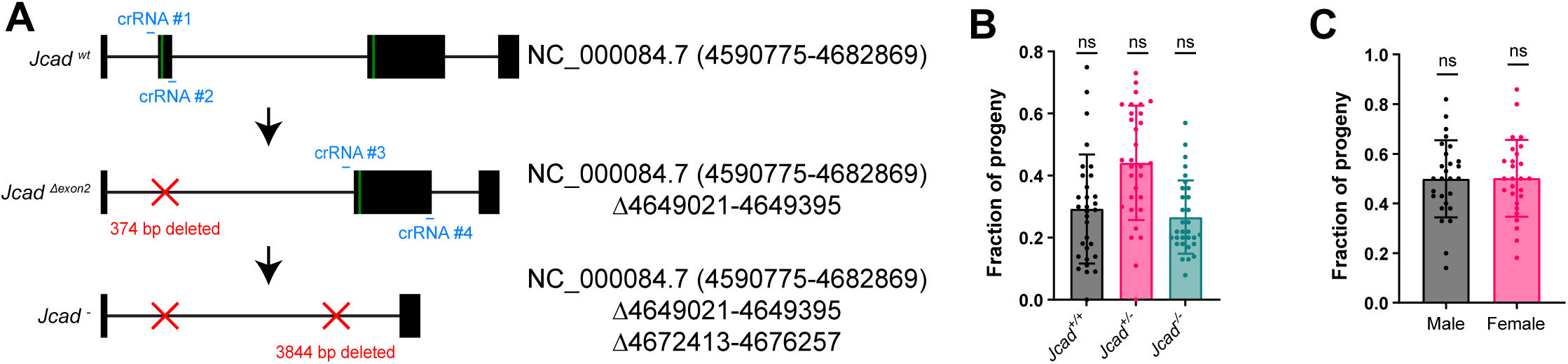
Generation of *Jcad* null allele using iGONAD. **(A)** Schematic of *Jcad* locus on GRCm39 C57BL/6J Chromosome 18 indicating loci deleted by iGONAD CRISPR/Cas9 genome editing. **(B)** Quantification of Mendelian ratios from *Jcad^+/-^*heterozygous intercrosses (N = 31 litters, one-sample *t* test). **(C)** Quantification of progeny sex from *Jcad^+/-^* heterozygous intercrosses (N = 27 litters, one-sample *t* test).

**Supplemental Figure S2:**
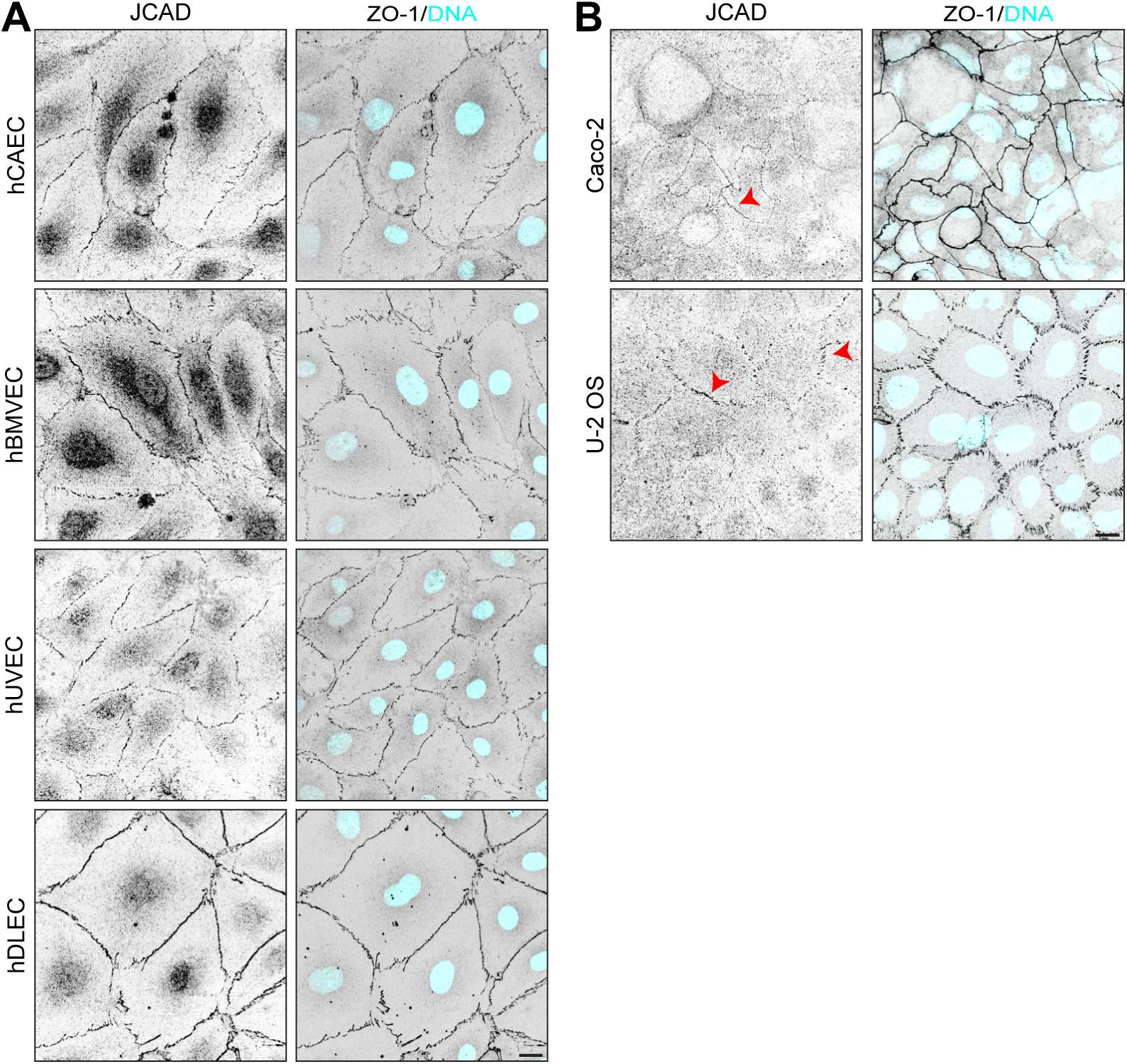
JCAD colocalizes with ZO-1 in epithelial cells and in endothelial cells of varying organotypic origin. **(A)** Immunostaining for JCAD and ZO-1 in human coronary artery endothelial cells (hCAECs), human brain microvascular endothelial cells (hBMVECs), human umbilical vein endothelial cells (hUVECs), and human dermal lymphatic endothelial cells (hDLECs). **(B)** Immunostaining for JCAD and ZO-1 in Caco-2 and U-2 OS cells. Arrowheads, examples of JCAD enriched tight junctions. Scale bars, 15 µm.

**Supplemental Figure S3:**
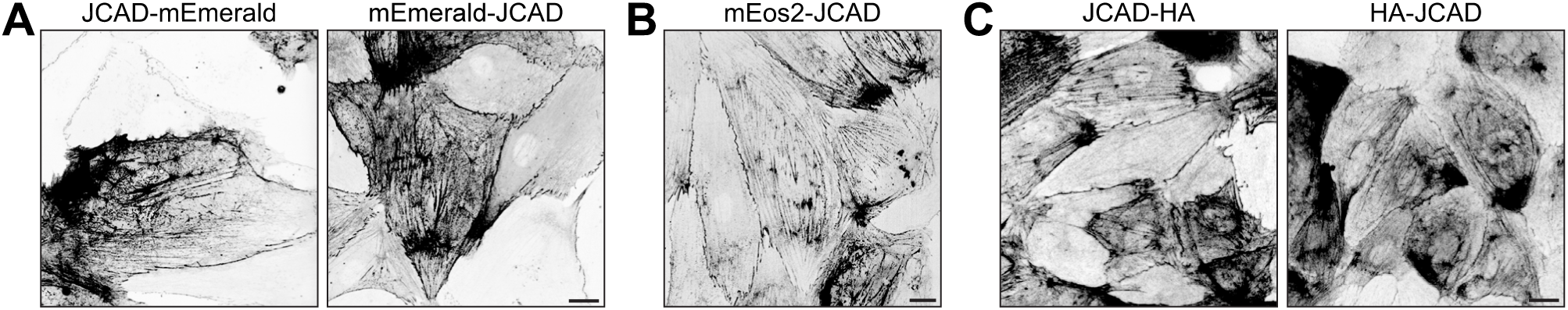
JCAD localization to filamentous network is independent of tagging. **(A)** Live-cell images of hMVECs expressing JCAD tagged with mEmerald at the N or C terminus and **(B)** mEos2 at the N terminus. **(C)** Immunostaining for the HA-tag in hMVECs expressing N- or C-terminally HA-tagged JCAD. Scale bars, 15 µm.

**Supplemental Figure S4:**
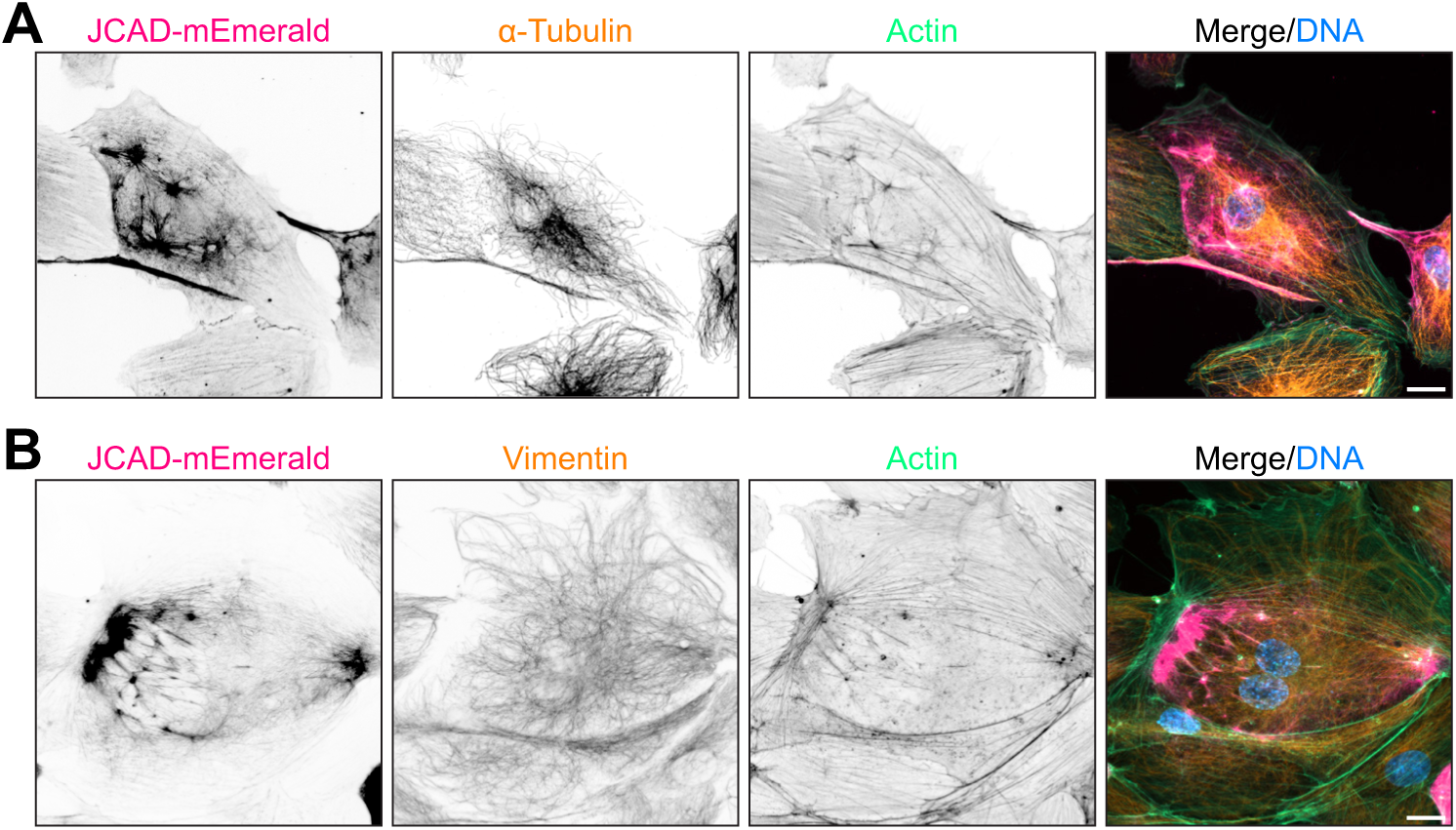
Overexpressed JCAD does not colocalize with intermediate filaments or microtubules. (A/B) Immunostaining for α-tubulin (A) and vimentin (B), with phalloidin-labeled F-actin, in hMVECs expressing JCAD-mEmerald. Scale bars, 15 µm.

**Supplemental Figure S5:**
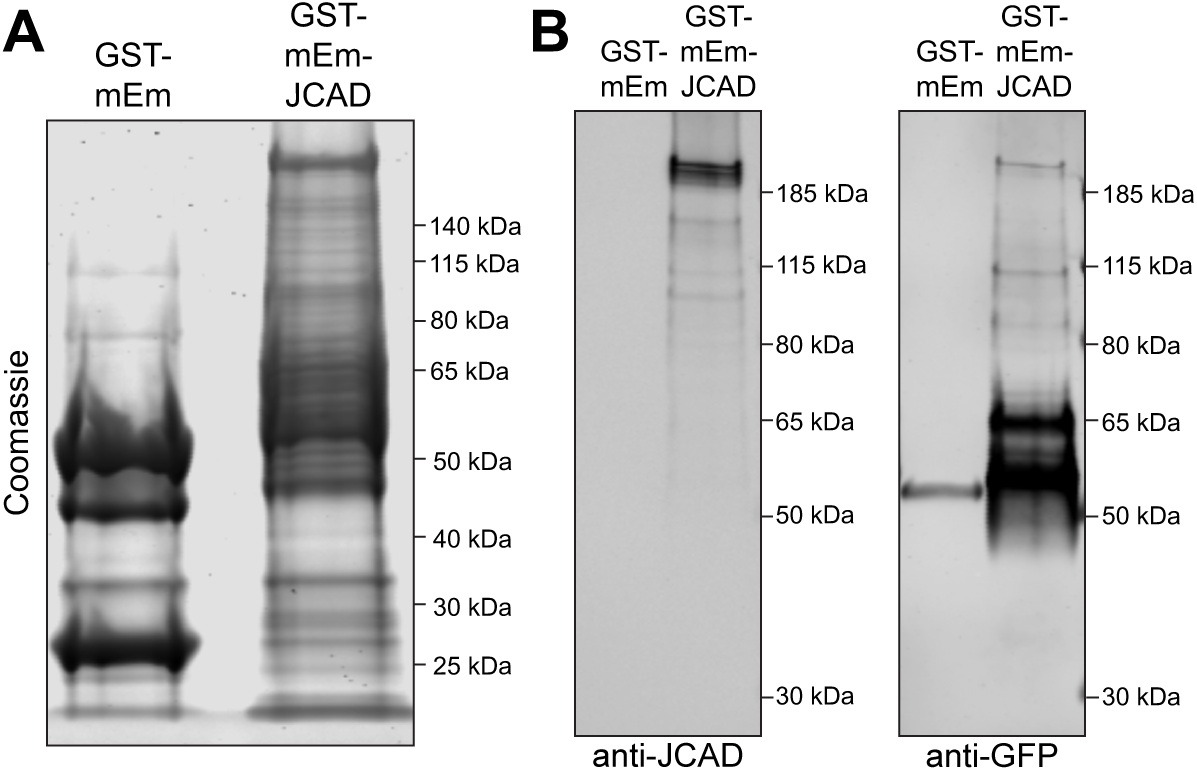
Validation of recombinant JCAD purification. **(A)** Coomassie-stained SDS-PAGE gel showing the purified GST-mEmerald and GST-mEmerald-JCAD fusion proteins after elution. **(B)** Western blot confirming the presence of bands at expected molecular weights, detected with a C-terminal JCAD antibody and anti-GFP antibody.

**Supplemental Figure S6:**
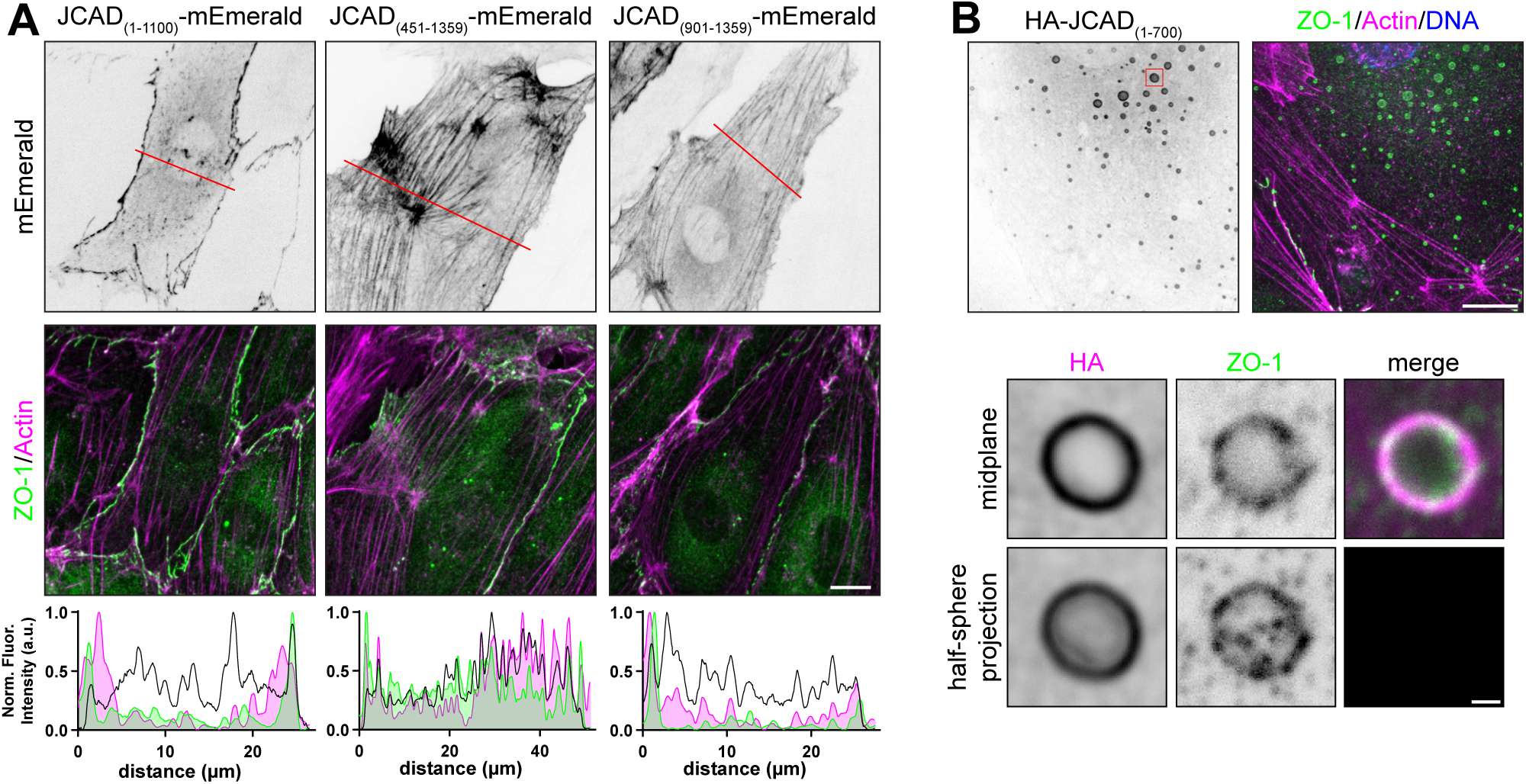
Additional JCAD truncations validate domain-specific functions. **(A)** Immunostaining for ZO-1 and actin in hMVECs expressing mEmerald or mEmerald-tagged JCAD truncations in a JCAD KD background. Red lines, location of displayed line-scan intensity profile. Scale bar, 10 µm. **(B)** Immunostaining for ZO-1 and actin in hMVECs expressing HA-JCAD_(1-700)_ exhibiting non-junctional aggregation. Scale bar, 10 µm. Red box, location of inset. Insets demonstrate **(i)** single confocal plane at middle of spherical droplet and **(ii)** a maximum intensity projection of confocal planes spanning bottom of droplet to mid-plane. Inset scale bar, 0.5 µm.

**Supplemental Figure S7:**
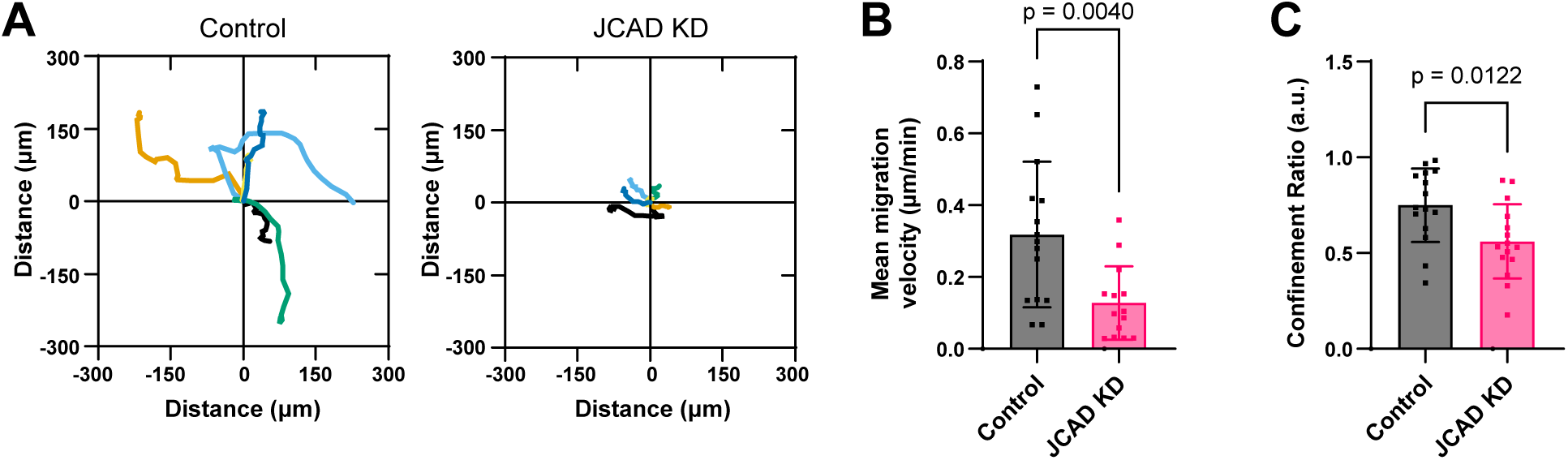
JCAD regulates random migration of hMVECs. **(A)** Representative cell migration tracks for Control and JCAD KD cells over 12 hours of live cell imaging. Each colored line represents the migratory path of a single cell**. (B/C)** Quantification of mean migration velocity and confinement ratio, defined as the distance traveled divided by displacement (N = 3 independent experiments; n = 4 - 6 tracks per N; Welch’s *t*-test).

## References

Abramson, J., J. Adler, J. Dunger, R. Evans, T. Green, A. Pritzel, O. Ronneberger, L. Willmore, A.J. Ballard, J. Bambrick, S.W. Bodenstein, D.A. Evans, C.-C. Hung, M. O’Neill, D. Reiman, K. Tunyasuvunakool, Z. Wu, A. Žemgulytė, E. Arvaniti, C. Beattie, O. Bertolli, A. Bridgland, A. Cherepanov, M. Congreve, A.I. Cowen-Rivers, A. Cowie, M. Figurnov, F.B. Fuchs, H. Gladman, R. Jain, Y.A. Khan, C.M.R. Low, K. Perlin, A. Potapenko, P. Savy, S. Singh, A. Stecula, A. Thillaisundaram, C. Tong, S. Yakneen, E.D. Zhong, M. Zielinski, A. Žídek, V. Bapst, P. Kohli, M. Jaderberg, D. Hassabis, and J.M. Jumper. 2024. Accurate structure prediction of biomolecular interactions with AlphaFold 3. Nature. 630:493–500. doi:10.1038/s41586-024-07487-w.

Akashi, M., T. Higashi, S. Masuda, T. Komori, and M. Furuse. 2011. A coronary artery disease-associated gene product, JCAD/KIAA1462, is a novel component of endothelial cell-cell junctions. Biochem Biophys Res Commun. 413:224–229. doi:10.1016/j.bbrc.2011.08.073.

Aw, W.Y., A. Sawhney, M. Rathod, C.P. Whitworth, E.L. Doherty, E. Madden, J. Lu, K. Westphal, R. Stack, and W.J. Polacheck. 2025. Dysfunctional mechanotransduction regulates the progression of PIK3CA-driven vascular malformations. APL Bioeng. 9:016106. doi:10.1063/5.0234507.

Barry, A.K., N. Wang, and D.E. Leckband. 2015. Local VE-cadherin mechanotransduction triggers long-ranged remodeling of endothelial monolayers. J Cell Sci. 128:1341–1351. doi:10.1242/jcs.159954.

Beutel, O., R. Maraspini, K. Pombo-García, C. Martin-Lemaitre, and A. Honigmann. 2019. Phase Separation of Zonula Occludens Proteins Drives Formation of Tight Junctions. Cell. 179:923–936.e11. doi:10.1016/j.cell.2019.10.011.

Boyd, J., B. Luo, S. Peri, B. Wirchansky, L. Hughes, C. Forsythe, and H. Wu. 2013. Whole exome sequence analysis of serous borderline tumors of the ovary. Gynecologic Oncology. 130:560–564. doi:10.1016/j.ygyno.2013.06.007.

Cao, J., and H. Schnittler. 2019. Putting VE-cadherin into JAIL for junction remodeling. Journal of Cell Science. 132:jcs222893. doi:10.1242/jcs.222893.

Carrithers, M., S. Tandon, S. Canosa, M. Michaud, D. Graesser, and J.A. Madri. 2005. Enhanced susceptibility to endotoxic shock and impaired STAT3 signaling in CD31-deficient mice. Am J Pathol. 166:185–196. doi:10.1016/S0002-9440(10)62243-2.

Citi, S., S. Paschoud, P. Pulimeno, F. Timolati, F. De Robertis, L. Jond, and L. Guillemot. 2009. The Tight Junction Protein Cingulin Regulates Gene Expression and RhoA Signaling. Annals of the New York Academy of Sciences. 1165:88–98. doi:10.1111/j.1749-6632.2009.04053.x.

Conway, D.E., M.T. Breckenridge, E. Hinde, E. Gratton, C.S. Chen, and M.A. Schwartz. 2013. Fluid shear stress on endothelial cells modulates mechanical tension across VE-cadherin and PECAM-1. Curr Biol. 23:1024–1030. doi:10.1016/j.cub.2013.04.049.

Corada, M., M. Mariotti, G. Thurston, K. Smith, R. Kunkel, M. Brockhaus, M.G. Lampugnani, I. Martin-Padura, A. Stoppacciaro, L. Ruco, D.M. McDonald, P.A. Ward, and E. Dejana. 1999. Vascular endothelial–cadherin is an important determinant of microvascular integrity in vivo. Proc. Natl. Acad. Sci. U.S.A. 96:9815–9820. doi:10.1073/pnas.96.17.9815.

Doube, M., M.M. Kłosowski, I. Arganda-Carreras, F.P. Cordelières, R.P. Dougherty, J.S. Jackson, B. Schmid, J.R. Hutchinson, and S.J. Shefelbine. 2010. BoneJ: Free and extensible bone image analysis in ImageJ. Bone. 47:1076–1079. doi:10.1016/j.bone.2010.08.023.

Douglas, G., V. Mehta, A. Al Haj Zen, I. Akoumianakis, A. Goel, V.S. Rashbrook, L. Trelfa, L. Donovan, E. Drydale, S. Chuaiphichai, C. Antoniades, H. Watkins, T. Kyriakou, E. Tzima, and K.M. Channon. 2020. A key role for the novel coronary artery disease gene JCAD in atherosclerosis via shear stress mechanotransduction. Cardiovascular Research. 116:1863–1874. doi:10.1093/cvr/cvz263.

Dunworth, W.P., K.L. Fritz-Six, and K.M. Caron. 2008. Adrenomedullin stabilizes the lymphatic endothelial barrier in vitro and in vivo. Peptides. 29:2243–2249. doi:10.1016/j.peptides.2008.09.009.

Eckenstaler, R., M. Hauke, and R.A. Benndorf. 2022. A current overview of RhoA, RhoB, and RhoC functions in vascular biology and pathology. Biochemical Pharmacology. 206:115321. doi:10.1016/j.bcp.2022.115321.

Efimova, N., and T.M. Svitkina. 2018. Branched actin networks push against each other at adherens junctions to maintain cell–cell adhesion. Journal of Cell Biology. 217:1827–1845. doi:10.1083/jcb.201708103.

Erdmann, J., C. Willenborg, J. Nahrstaedt, M. Preuss, I.R. König, J. Baumert, P. Linsel-Nitschke, C. Gieger, S. Tennstedt, P. Belcredi, Z. Aherrahrou, N. Klopp, C. Loley, K. Stark, C. Hengstenberg, P. Bruse, J. Freyer, A.K. Wagner, A. Medack, W. Lieb, A. Großhennig, H.B. Sager, A. Reinhardt, A. Schäfer, S. Schreiber, N.E. El Mokhtari, D. Raaz-Schrauder, T. Illig, C.D. Garlichs, A.B. Ekici, A. Reis, J. Schrezenmeir, D. Rubin, A. Ziegler, H.E. Wichmann, A. Doering, C. Meisinger, T. Meitinger, A. Peters, and H. Schunkert. 2011. Genome-wide association study identifies a new locus for coronary artery disease on chromosome 10p11.23. European Heart Journal. 32:158–168. doi:10.1093/eurheartj/ehq405.

Giampietro, C., A. Disanza, L. Bravi, M. Barrios-Rodiles, M. Corada, E. Frittoli, C. Savorani, M.G. Lampugnani, B. Boggetti, C. Niessen, J.L. Wrana, G. Scita, and E. Dejana. 2015. The actin-binding protein EPS8 binds VE-cadherin and modulates YAP localization and signaling. J Cell Biol. 211:1177–1192. doi:10.1083/jcb.201501089.

Gimbrone, M.A., and G. García-Cardeña. 2016. Endothelial Cell Dysfunction and the Pathobiology of Atherosclerosis. Circulation Research. 118:620–636. doi:10.1161/CIRCRESAHA.115.306301.

Grazia Lampugnani, M., A. Zanetti, M. Corada, T. Takahashi, G. Balconi, F. Breviario, F. Orsenigo, A. Cattelino, R. Kemler, T.O. Daniel, and E. Dejana. 2003. Contact inhibition of VEGF-induced proliferation requires vascular endothelial cadherin, beta-catenin, and the phosphatase DEP-1/CD148. J Cell Biol. 161:793–804. doi:10.1083/jcb.200209019.

Grimsley-Myers, C.M., R.H. Isaacson, C.M. Cadwell, J. Campos, M.S. Hernandes, K.R. Myers, T. Seo, W. Giang, K.K. Griendling, and A.P. Kowalczyk. 2020. VE-cadherin endocytosis controls vascular integrity and patterning during development. J Cell Biol. 219:e201909081. doi:10.1083/jcb.201909081.

Gurumurthy, C.B., M. Sato, A. Nakamura, M. Inui, N. Kawano, M.A. Islam, S. Ogiwara, S. Takabayashi, M. Matsuyama, S. Nakagawa, H. Miura, and M. Ohtsuka. 2019. Creation of CRISPR-based germline-genome-engineered mice without ex vivo handling of zygotes by i-GONAD. Nat Protoc. 14:2452–2482. doi:10.1038/s41596-019-0187-x.

Hahn, C., and M.A. Schwartz. 2009. Mechanotransduction in vascular physiology and atherogenesis. Nat Rev Mol Cell Biol. 10:53–62. doi:10.1038/nrm2596.

Hara, T., T. Monguchi, N. Iwamoto, M. Akashi, K. Mori, T. Oshita, M. Okano, R. Toh, Y. Irino, M. Shinohara, Y. Yamashita, G. Shioi, M. Furuse, T. Ishida, and K.I. Hirata. 2017. Targeted Disruption of JCAD (Junctional Protein Associated with Coronary Artery Disease)/KIAA1462, a Coronary Artery Disease-Associated Gene Product, Inhibits Angiogenic Processes in Vitro and in Vivo. Arteriosclerosis, Thrombosis, and Vascular Biology. 37:1667–1673. doi:10.1161/ATVBAHA.117.309721.

Heuzé, M.L., G.H.N. Sankara Narayana, J. D’Alessandro, V. Cellerin, T. Dang, D.S. Williams, J.C. Van Hest, P. Marcq, R.-M. Mège, and B. Ladoux. 2019. Myosin II isoforms play distinct roles in adherens junction biogenesis. eLife. 8:e46599. doi:10.7554/eLife.46599.

Hiraoka, Y., M. Matsumura, Y. Kakei, D. Takeda, M. Shigeoka, A. Kimoto, T. Hasegawa, and M. Akashi. 2023. Expression of JCAD and EGFR in Perineurial Cell-Cell Junctions of Human Inferior Alveolar Nerve. J Histochem Cytochem. 71:321–332. doi:10.1369/00221554231182193.

Jones, P.D., M.A. Kaiser, M.G. Najafabadi, S. Koplev, Y. Zhao, G. Douglas, T. Kyriakou, S. Andrews, R. Rajmohan, H. Watkins, K.M. Channon, S. Ye, X. Yang, J.L.M. Björkegren, N.J. Samani, and T.R. Webb. 2018. JCAD, a Gene at the 10p11 Coronary Artery Disease Locus, Regulates Hippo Signaling in Endothelial Cells. Arteriosclerosis, Thrombosis, and Vascular Biology. 38:1711–1722. doi:10.1161/ATVBAHA.118.310976.

Kim, D.I., B. Kc, W. Zhu, K. Motamedchaboki, V. Doye, and K.J. Roux. 2014. Probing nuclear pore complex architecture with proximity-dependent biotinylation. Proc. Natl. Acad. Sci. U.S.A. 111:E2453–E2461. doi:10.1073/pnas.1406459111.

Koyama, T., L. Ochoa-Callejero, T. Sakurai, A. Kamiyoshi, Y. Ichikawa-Shindo, N. Iinuma, T. Arai, T. Yoshizawa, Y. Iesato, Y. Lei, R. Uetake, A. Okimura, A. Yamauchi, M. Tanaka, K. Igarashi, Y. Toriyama, H. Kawate, R.H. Adams, H. Kawakami, N. Mochizuki, A. Martínez, and T. Shindo. 2013. Vascular Endothelial Adrenomedullin-RAMP2 System Is Essential for Vascular Integrity and Organ Homeostasis. Circulation. 127:842–853. doi:10.1161/CIRCULATIONAHA.112.000756.

Lampugnani, M.G., F. Orsenigo, M.C. Gagliani, C. Tacchetti, and E. Dejana. 2006. Vascular endothelial cadherin controls VEGFR-2 internalization and signaling from intracellular compartments. J Cell Biol. 174:593–604. doi:10.1083/jcb.200602080.

Lampugnani, M.G., M. Resnati, M. Raiteri, R. Pigott, A. Pisacane, G. Houen, L.P. Ruco, and E. Dejana. 1992. A novel endothelial-specific membrane protein is a marker of cell-cell contacts. Journal of Cell Biology. 118:1511–1522. doi:10.1083/jcb.118.6.1511.

Landau, L.M., N. Chaudhary, Y.C. Tien, M. Rogozinska, S. Joshi, C. Yao, J. Crowley, K. Hullahalli, I.W. Campbell, M.K. Waldor, M. Haigis, and J.C. Kagan. 2024. pLxIS-containing domains are biochemically flexible regulators of interferons and metabolism. Mol Cell. 84:2436–2454.e10. doi:10.1016/j.molcel.2024.05.030.

Libby, P. 2002. Inflammation in atherosclerosis. Nature. 420:868–874. doi:10.1038/nature01323.

Liberale, L., Y.M. Puspitasari, S. Ministrini, A. Akhmedov, S. Kraler, N.R. Bonetti, G. Beer, A. Vukolic, D. Bongiovanni, J. Han, K. Kirmes, I. Bernlochner, J. Pelisek, J.H. Beer, Z.-G. Jin, D. Pedicino, G. Liuzzo, K. Stellos, F. Montecucco, F. Crea, T.F. Lüscher, and G.G. Camici. 2023. JCAD promotes arterial thrombosis through PI3K/Akt modulation: a translational study. European Heart Journal. 44:1818–1833. doi:10.1093/eurheartj/ehac641.

Lin, Z., Z. Yang, R. Xie, Z. Ji, K. Guan, and M. Zhang. 2019. Decoding WW domain tandem-mediated target recognitions in tissue growth and cell polarity. eLife. 8:e49439. doi:10.7554/eLife.49439.

Lindner, V., and H. Heinle. 1982. Binding properties of circulating Evans blue in rabbits as determined by disc electrophoresis. Atherosclerosis. 43:417–422. doi:10.1016/0021-9150(82)90040-5.

Maddala, R., and P.V. Rao. 2020. Global phosphotyrosinylated protein profile of cell-matrix adhesion complexes of trabecular meshwork cells. American Journal of Physiology - Cell Physiology. 319:C288–C299. doi:10.1152/ajpcell.00537.2019.

Mayo, L.N., F. Duong, A. Mompeón, K.A. Jacobs, M.L. Iruela-Arispe, and M.L. Kutys. 2026. Scrib organizes cortical actomyosin clusters to maintain adherens junctions and angiogenic sprouting. Journal of Cell Biology. 225:e202503146. doi:10.1083/jcb.202503146.

Montagne, A., Z. Zhao, and B.V. Zlokovic. 2017. Alzheimer’s disease: A matter of blood-brain barrier dysfunction? J Exp Med. 214:3151–3169. doi:10.1084/jem.20171406.

Morini, M.F., C. Giampietro, M. Corada, F. Pisati, E. Lavarone, S.I. Cunha, L.L. Conze, N. O’Reilly, D. Joshi, S. Kjaer, R. George, E. Nye, A. Ma, J. Jin, R. Mitter, M. Lupia, U. Cavallaro, D. Pasini, D.P. Calado, E. Dejana, and A. Taddei. 2018. VE-Cadherin–Mediated Epigenetic Regulation of Endothelial Gene Expression. Circulation Research. 122:231–245. doi:10.1161/CIRCRESAHA.117.312392.

Muller, W.A. 2003. Leukocyte-endothelial-cell interactions in leukocyte transmigration and the inflammatory response. Trends Immunol. 24:327–334. doi:10.1016/s1471-4906(03)00117-0.

Murdock, D.G., Y. Bradford, N. Schnetz-Boutaud, P. Mayo, M.J. Allen, L.N. D’Aoust, X. Liang, S.L. Mitchell, S. Zuchner, G.W. Small, J.R. Gilbert, M.A. Pericak-Vance, and J.L. Haines. 2013. KIAA1462, a coronary artery disease associated gene, is a candidate gene for late onset Alzheimer disease in APOE carriers. PLoS ONE. 8:1–9. doi:10.1371/journal.pone.0082194.

Ohnishi, H., T. Nakahara, K. Furuse, H. Sasaki, S. Tsukita, and M. Furuse. 2004. JACOP, a novel plaque protein localizing at the apical junctional complex with sequence similarity to cingulin. J Biol Chem. 279:46014–46022. doi:10.1074/jbc.M402616200.

Ohtsuka, M., M. Sato, H. Miura, S. Takabayashi, M. Matsuyama, T. Koyano, N. Arifin, S. Nakamura, K. Wada, and C.B. Gurumurthy. 2018. i-GONAD: a robust method for in situ germline genome engineering using CRISPR nucleases. Genome Biol. 19:25. doi:10.1186/s13059-018-1400-x.

Peden, J.F., J.C. Hopewell, D. Saleheen, J.C. Chambers, J. Hager, N. Soranzo, R. Collins, J. Danesh, P. Elliott, M. Farrall, K. Stirrups, W. Zhang, A. Hamsten, S. Parish, M. Lathrop, H. Watkins, R. Clarke, P. Deloukas, J.S. Kooner, A. Goel, H. Ongen, R.J. Strawbridge, S. Heath, A. MaΪarstig, A. Helgadottir, J. Öhrvik, M. Murtaza, S. Potter, S.E. Hunt, M. Delepine, S. Jalilzadeh, T. Axelsson, A.C. Syvanen, R. Gwilliam, S. Bumpstead, E. Gray, S. Edkins, L. Folkersen, T. Kyriakou, A. Franco-Cereceda, A. Gabrielsen, U. Seedorf, P. Eriksson, A. Offer, L. Bowman, P. Sleight, J. Armitage, R. Peto, G. Abecasis, N. Ahmed, M. Caulfield, P. Donnelly, P. Froguel, A.S. Kooner, M.I. McCarthy, N.J. Samani, J. Scott, J. Sehmi, A. Silveira, M.L. Hellánius, F.M. Van ’T Hooft, G. Olsson, S. Rust, G. Assmann, S. Barlera, G. Tognoni, M.G. Franzosi, P. Linksted, F.R. Green, A. Rasheed, M. Zaidi, N. Shah, M. Samuel, N.H. Mallick, M. Azhar, K.S. Zaman, A. Samad, M. Ishaq, A.R. Gardezi, F.U.R. Memon, P.M. Frossard, T. Spector, L. Peltonen, M.S. Nieminen, J. Sinisalo, V. Salomaa, S. Ripatti, D. Bennett, K. Leander, B. Gigante, U. De Faire, S. Pietri, F. Gori, R. Marchioli, S. Sivapalaratnam, J.J.P. Kastelein, M.D. Trip, E.V. Theodoraki, et al. 2011. A genome-wide association study in Europeans and South Asians identifies five new loci for coronary artery disease. Nature Genetics. 43:339–346. doi:10.1038/ng.782.

Polacheck, W.J., M.L. Kutys, J. Yang, J. Eyckmans, Y. Wu, H. Vasavada, K.K. Hirschi, and C.S. Chen. 2017. A non-canonical Notch complex regulates adherens junctions and vascular barrier function. Nature. 552:258–262. doi:10.1038/nature24998.

Pombo-García, K., O. Adame-Arana, C. Martin-Lemaitre, F. Jülicher, and A. Honigmann. 2024. Membrane prewetting by condensates promotes tight-junction belt formation. Nature. 632:647–655. doi:10.1038/s41586-024-07726-0.

Radu, M., and J. Chernoff. 2013. An in vivo Assay to Test Blood Vessel Permeability. JoVE. 50062. doi:10.3791/50062.

Rathod, M.L., W.Y. Aw, S. Huang, J. Lu, E.L. Doherty, C.P. Whithworth, G. Xi, P. Roy-Chaudhury, and W.J. Polacheck. 2024. Donor-Derived Engineered Microvessels for Cardiovascular Risk Stratification of Patients with Kidney Failure. Small. 20:e2307901. doi:10.1002/smll.202307901.

Ross, R., and J.A. Glomset. 1976. The Pathogenesis of Atherosclerosis. N Engl J Med. 295:369–377. doi:10.1056/NEJM197608122950707.

Rouaud, F., W. Huang, A. Flinois, K. Jain, E. Vasileva, T. Di Mattia, M. Mauperin, D.A.D. Parry, V. Dugina, C. Chaponnier, I. Méan, S. Montessuit, A. Mutero-Maeda, J. Yan, and S. Citi. 2023. Cingulin and paracingulin tether myosins-2 to junctions to mechanoregulate the plasma membrane. J Cell Biol. 222:e202208065. doi:10.1083/jcb.202208065.

Rubinson, D.A., C.P. Dillon, A.V. Kwiatkowski, C. Sievers, L. Yang, J. Kopinja, D.L. Rooney, M. Zhang, M.M. Ihrig, M.T. McManus, F.B. Gertler, M.L. Scott, and L. Van Parijs. 2003. A lentivirus-based system to functionally silence genes in primary mammalian cells, stem cells and transgenic mice by RNA interference. Nat Genet. 33:401–406. doi:10.1038/ng1117.

Schindelin, J., I. Arganda-Carreras, E. Frise, V. Kaynig, M. Longair, T. Pietzsch, S. Preibisch, C. Rueden, S. Saalfeld, B. Schmid, J.-Y. Tinevez, D.J. White, V. Hartenstein, K. Eliceiri, P. Tomancak, and A. Cardona. 2012. Fiji: an open-source platform for biological-image analysis. Nat Methods. 9:676–682. doi:10.1038/nmeth.2019.

Schwayer, C., S. Shamipour, K. Pranjic-Ferscha, A. Schauer, M. Balda, M. Tada, K. Matter, and C.-P. Heisenberg. 2019. Mechanosensation of Tight Junctions Depends on ZO-1 Phase Separation and Flow. Cell. 179:937–952.e18. doi:10.1016/j.cell.2019.10.006.

Senger, D.R., S.J. Galli, A.M. Dvorak, C.A. Perruzzi, V.S. Harvey, and H.F. Dvorak. 1983. Tumor cells secrete a vascular permeability factor that promotes accumulation of ascites fluid. Science. 219:983–985. doi:10.1126/science.6823562.

Serafin, D.S., N.R. Harris, L. Bálint, E.S. Douglas, and K.M. Caron. 2024. Proximity interactome of lymphatic VE-cadherin reveals mechanisms of junctional remodeling and reelin secretion. Nat Commun. 15:7734. doi:10.1038/s41467-024-51918-1.

Simons, M. 2023. Endothelial-to-mesenchymal transition: advances and controversies. Curr Opin Physiol. 34:100678. doi:10.1016/j.cophys.2023.100678.

Singh, T., K.A. Jacobs, W.J. Polacheck, and M.L. Kutys. 2026. The Notch1 intracellular domain orchestrates mechanotransduction of fluid shear stress. Life Sci. Alliance. 9:e202503599. doi:10.26508/lsa.202503599.

Stehbens, S., H. Pemble, L. Murrow, and T. Wittmann. 2012. Imaging intracellular protein dynamics by spinning disk confocal microscopy. Methods Enzymol. 504:293–313. doi:10.1016/B978-0-12-391857-4.00015-X.

Stephenson, R.E., T. Higashi, I.S. Erofeev, T.R. Arnold, M. Leda, A.B. Goryachev, and A.L. Miller. 2019. Rho Flares Repair Local Tight Junction Leaks. Dev Cell. 48:445–459.e5. doi:10.1016/j.devcel.2019.01.016.

Sun, D., X. Zhao, T. Wiegand, C. Martin-Lemaitre, T. Borianne, L. Kleinschmidt, S.W. Grill, A.A. Hyman, C. Weber, and A. Honigmann. 2025. Assembly of tight junction belts by ZO1 surface condensation and local actin polymerization. Developmental Cell. 60:1234–1250.e6. doi:10.1016/j.devcel.2024.12.012.

Sweeney, M.D., A.P. Sagare, and B.V. Zlokovic. 2018. Blood-brain barrier breakdown in Alzheimer disease and other neurodegenerative disorders. Nat Rev Neurol. 14:133–150. doi:10.1038/nrneurol.2017.188.

Taddei, A., C. Giampietro, A. Conti, F. Orsenigo, F. Breviario, V. Pirazzoli, M. Potente, C. Daly, S. Dimmeler, and E. Dejana. 2008. Endothelial adherens junctions control tight junctions by VE-cadherin-mediated upregulation of claudin-5. Nat Cell Biol. 10:923–934. doi:10.1038/ncb1752.

Takeichi, M. 1977. Functional correlation between cell adhesive properties and some cell surface proteins. J Cell Biol. 75:464–474. doi:10.1083/jcb.75.2.464.

Tual-Chalot, S., K.R. Allinson, M. Fruttiger, and H.M. Arthur. 2013. Whole mount immunofluorescent staining of the neonatal mouse retina to investigate angiogenesis in vivo. J Vis Exp. e50546. doi:10.3791/50546.

Uhlen, M., C. Zhang, S. Lee, E. Sjöstedt, L. Fagerberg, G. Bidkhori, R. Benfeitas, M. Arif, Z. Liu, F. Edfors, K. Sanli, K. von Feilitzen, P. Oksvold, E. Lundberg, S. Hober, P. Nilsson, J. Mattsson, J.M. Schwenk, H. Brunnström, B. Glimelius, T. Sjöblom, P.-H. Edqvist, D. Djureinovic, P. Micke, C. Lindskog, A. Mardinoglu, and F. Ponten. 2017. A pathology atlas of the human cancer transcriptome. Science. 357:eaan2507. doi:10.1126/science.aan2507.

Van Den Biggelaar, M., J.R. Hernández-Fernaud, B.L. Van Den Eshof, L.J. Neilson, A.B. Meijer, K. Mertens, and S. Zanivan. 2014. Quantitative phosphoproteomics unveils temporal dynamics of thrombin signaling in human endothelial cells. Blood. 123:e22–e36. doi:10.1182/blood-2013-12-546036.

Van Itallie, C.M., A. Aponte, A.J. Tietgens, M. Gucek, K. Fredriksson, and J.M. Anderson. 2013. The N and C termini of ZO-1 are surrounded by distinct proteins and functional protein networks. Journal of Biological Chemistry. 288:13775–13788. doi:10.1074/jbc.M113.466193.

Vicente-Manzanares, M., X. Ma, R.S. Adelstein, and A.R. Horwitz. 2009. Non-muscle myosin II takes centre stage in cell adhesion and migration. Nat Rev Mol Cell Biol. 10:778–790. doi:10.1038/nrm2786.

Weißenbruch, K., and R. Mayor. 2024. Actomyosin forces in cell migration: Moving beyond cell body retraction. BioEssays. 46:2400055. doi:10.1002/bies.202400055.

Wildenberg, G.A., M.R. Dohn, R.H. Carnahan, M.A. Davis, N.A. Lobdell, J. Settleman, and A.B. Reynolds. 2006. p120-Catenin and p190RhoGAP Regulate Cell-Cell Adhesion by Coordinating Antagonism between Rac and Rho. Cell. 127:1027–1039. doi:10.1016/j.cell.2006.09.046.

Xia, J., W.Y. Wang, K.A. Jacobs, D. Lin, E.H. Jarman, D.L. Matera, K. Loesel, C.D. Davidson, H.L. Hiraki, X. Tan, E.H. Shikanov, R.N. Kent, C. Parent, X. Fan, A. Shikanov, M.L. Kutys, and B.M. Baker. 2025. Perivascular matrix densification dysregulates angiogenesis and activates pro-inflammatory endothelial cells. doi:10.1101/2025.05.20.655215.

Xu, S., Y. Xu, P. Liu, S. Zhang, H. Liu, S. Slavin, S. Kumar, M. Koroleva, J. Luo, X. Wu, A. Rahman, J. Pelisek, H. Jo, S. Si, C.L. Miller, and Z.G. Jin. 2019. The novel coronary artery disease risk gene JCAD/KIAA1462 promotes endothelial dysfunction and atherosclerosis. European Heart Journal. 40:2398–2408. doi:10.1093/eurheartj/ehz303.

Xu, X.-P., S. Pokutta, M. Torres, M.F. Swift, D. Hanein, N. Volkmann, and W.I. Weis. 2020. Structural basis of αE-catenin–F-actin catch bond behavior. eLife. 9:e60878. doi:10.7554/eLife.60878.

Ye, J., T.S. Li, G. Xu, Y.M. Zhao, N.P. Zhang, J. Fan, and J. Wu. 2017. JCAD promotes progression of nonalcoholic steatohepatitis to liver cancer by inhibiting LATS2 kinase activity. Cancer Research. 77:5287–5300. doi:10.1158/0008-5472.CAN-17-0229.

Yuan, M., C. Zhang, K. Von Feilitzen, M. Zwahlen, M. Shi, X. Li, H. Yang, X. Song, H. Turkez, M. Uhlén, and A. Mardinoglu. 2025. The Human Pathology Atlas for deciphering the prognostic features of human cancers. EBioMedicine. 111:105495. doi:10.1016/j.ebiom.2024.105495.

Zlokovic, B.V. 2011. Neurovascular pathways to neurodegeneration in Alzheimer’s disease and other disorders. Nat Rev Neurosci. 12:723–738. doi:10.1038/nrn3114.

