## Supplemental Tables and Videos for "JCAD couples tight junction condensates to actin and RhoA to maintain the endothelial barrier": TableS1.pdf

**Table S1**

| Antibody | Source | Catalog # | Details |
|---|---|---|---|
| rabbit anti-KIAA1462 (JCAD) | Atlas Antibodies | HPA017956 | IF: 0.3-0.5 μg/ml; WB: 0.1 μg/ml |
| Rat anti-mouse CD31 | BD Biosciences | 553370 | IF: 1 μg/ml |
| mouse anti-human CD144 (V55-7H1) FITC | BD Biosciences | 560874 | IF: 1:50 |
| mouse anti-human CD144 (V55-7H1) Alexa Fluor 647 | BD Biosciences | 561567 | IF: 1:200 |
| rabbit anti-LARG (ARHGEF12) | Bethyl Laboratories | A301-960A | WB: 0.5 μg/ml |
| Rabbit anti-GAPDH (14C10) | Cell Signaling Technologies | 2118 | WB: 1:2000 |
| Rabbit anti-HA tag (C29F4) | Cell Signaling Technologies | 3724 | IF: 1:300; WB: 1:1000 |
| rabbit anti-vimentin | Cell Signaling Technologies | 3932 | IF: 1:300 |
| Anti-GFP, NeuroMab clone N86/8 | Developmental Studies Hybridoma Bank | N86/8 | WB: 0.5 μg/ml |
| Mouse anti-ZO-1 (ZO1-1A12) | Invitrogen | 33-9100 | IF: 2.5 μg/ml |
| Goat anti-Mouse IgG (H+L), Alexa Fluor 488 | Invitrogen | A11001 | IF: 4 μg/ml |
| Goat anti-Mouse IgG (H+L) , Alexa Fluor 647 | Invitrogen | A-21236 | IF: 4 μg/ml |
| Goat anti-Rabbit IgG (H+L), Alexa Fluor 647 | Invitrogen | A-21244 | IF: 4 μg/ml |
| Goat anti-Rabbit IgG (H+L), Alexa Fluor 568 | Invitrogen | A-11011 | IF: 4 μg/ml |
| Donkey anti-rat IgG Alexa Fluor 647 | Invitrogen | A-78947 | IF: 4 μg/ml |
| IRDye® 680RD Donkey Anti-Mouse IgG | LICORbio | 926-68072 | WB: 0.1 μg/ml |
| IRDye® 800CW Donkey Anti-Rabbit IgG | LICORbio | 926-32213 | WB: 0.1 μg/ml |
| mouse anti-HA-tag | Santa Cruz Biotechnology | sc-7392 | IF: 2-4 μg/ml; WB: 0.2 μg/ml |
| Mouse anti-JCAD (F-7) Alexa Fluor 488 | Santa Cruz Biotechnology | sc-515169 AF488 | IF: 2-4 μg/ml |
| Mouse anti-JCAD (F-7) | Santa Cruz Biotechnology | sc-515169 | WB: 2 μg/ml |
| Mouse anti-beta actin (C4) Alexa Fluor 680 | Santa Cruz Biotechnology | sc-47778 AF680 | WB: 0.2 μg/ml |
| mouse anti-VE-cadherin (F-8) | Santa Cruz Biotechnology | sc-9989 | IF: 1 μg/ml |
| mouse anti-α-tubulin (DM1A) | Sigma Aldrich | T9026 | IF: 1:200 |
| **Reagent** | **Source** | **Catalog #** | |
| Actin Binding Protein Spin-Down Assay Biochem Kit | Cytoskeleton Inc | bk013 | |
| Bovine Serum Albumin, Heat Shock Treated | Fisher Scientific | BP1600-100 | |
| Collagen I, Rat | Corning | 354236 | |
| D-(+)-Biotin | VWR | 97061 | |
| DAPI (4′,6-diamidino-2-phenylindole) | Thermo Scientific | D1306 | |
| Doxycycline hyclate | Sigma Aldrich | D9891 | |
| Dynabeads Streptavidin Magnetic Beads C1 | Invitrogen | 65001 | |
| Evans blue | Sigma Aldrich | E2129 | |
| EZ-Link Sulfo-NHS-LC-Biotin | Thermo Scientific | A39257 | |
| Fibronectin, Human | Corning | 354008 | |
| Halt Protease and Phosphatase Inhibitor Cocktail, EDTA-free (100X) | Thermo Scientific | 78443 | |
| Halt Protease Inhibitor Cocktail, EDTA-free (100X) | Thermo Scientific | 78425 | |
| Isopropyl-B-D-thiogalactopyranoside | Zymo Research | 11-230 | |
| Latrunculin A | Cayman Chemical | 10010630-100 | |
| Lipopolysaccharides from Escherichia coli O111:B4 | Sigma Aldrich | L4391 | |
| Normal Goat Serum | Jackson Immuno Research Labs | 5000121 | |

| | | |
|---|---|---|
| rhodamine phalloidin | Invitrogen | R415 |
| SPY555-Fast act | Cytoskeleton Inc | SC205 |
| Streptavidin, Alexa Fluor 568 | Invitrogen | S11226 |
| Thrombin, Bovine, High Activity | Sigma Aldrich | 604980 |

| **Plasmid** | **Source** |
|---|---|
| pRRL VE-cadherin-BirA* | Mayo, et al., J Cell Bio, 2026 |
| pLentiCRISPRv2 Scramble sgRNA | Polacheck, et al., Nature, 2017 |
| pLentiCRISPRv2 CDH5 sgRNA | Polacheck, et al., Nature, 2017 |
| pLL3.7 PuroR NS shRNA | This study |
| pLL3.7 PuroR JCAD shRNA | This study |
| pRRL JCAD-mEmerald | This study |
| pRRL mEmerald-JCAD | This study |
| pRRL VE-cadherin-mApple | Addgene Plasmid #54959 |
| pRRL mScarlet-ZO-1 | This study |
| pRRL mEmerald | This study |
| pCW57 JCAD-mEmerald | This study |
| pGEX GST-mEmerald | This study |
| pGEX GST-mEmerald-JCAD | This study |
| pRRL lifeact-mScarlet-P2A-HygroR | This study |
| pLL3.7 JCAD shNRA PuroR-IRES2-mEmerald | This study |
| pLL3.7 JCAD shNRA PuroR-IRES2-JCAD-mEmerald | This study |
| pLL3.7 JCAD shNRA PuroR-IRES2-JCAD(1-450)-mEmerald | This study |
| pLL3.7 JCAD shNRA PuroR-IRES2-JCAD(1-900)-mEmerald | This study |
| pLL3.7 JCAD shNRA PuroR-IRES2-JCAD(1-1100)-mEmerald | This study |
| pLL3.7 JCAD shNRA PuroR-IRES2-JCAD(451-1359)-mEmerald | This study |
| pLL3.7 JCAD shNRA PuroR-IRES2-JCAD(901-1359)-mEmerald | This study |
| pLL3.7 JCAD shNRA PuroR-IRES2-JCAD(1101-1359)-mEmerald | This study |
| pRRL mEos2-JCAD | This study |
| pRRL JCAD-HA | This study |
| pRRL HA-JCAD | This study |
| pRRL HA-JCAD$_{(1-700)}$ | This study |
| pRRL HA-JCAD$_{(1-450)}$ | This study |
| pRRL HA-JCAD$_{(451-700)}$ | This study |
| pGEX GST-6P-2 | A gift from Diane Barber |
| pGEX GST-RhoA | This study |
| pGEX GST-RhoA G17A | This study |
| pGEX GST-RhoA T19N | This study |
| pGEX-4T3-RhoA-Q63L | Addgene Plasmid #12961 |
