## Supplemental Tables and Videos for "JCAD couples tight junction condensates to actin and RhoA to maintain the endothelial barrier": TableS2.pdf

**Table S2**

| Primer name | Description | Sequence |
|---|---|---|
| Jcad-ex2-WT_fwd | genotyping of exon 2 deletion allele | TGAGGTTTGATGACGCTGC |
| Jcad-ex2-mut_fwd | | TGGGTGTCCACTGAGGCACG |
| Jcad-ex2_rev | | CAACATCGGTGCAAAGGCAA |
| Jcad-ex3_fwd | genotyping of exon 3 deletion, to identify mice carrying Jcad null allele | GACTGCAGGGAGAATATTCC |
| Jcad-ex3-mut_rev | | GAGGTCTGAGTTGTTTATAAGG |
| Jcad-ex3-WT_rev | | CTGTAAGCCTGAGCCATTCC |
