## Supplemental Tables and Videos for "JCAD couples tight junction condensates to actin and RhoA to maintain the endothelial barrier": TableS3.pdf

**Table S3**

| Plasmid name | Description | Vector | Vector restriction enzymes or PCR primers | Insert | Primer 1 | Primer 2 |
|---|---|---|---|---|---|---|
| pLL3.7 PuroR | Lentiviral transfer plasmid for U6 promoter driven shRNA expression with puromycin resistance casette | pLL3.7 | fwd: CGTCGAGGGACCTAATAAC rev: TCTGACGGTTCACTAAAC | PuroR | tggtttagtgaaccgtcagaatgaccgagtacaagcccacgg | agttattaggtccctcgacgaattctcaggcaccgggcttgcg |
| pLL3.7 PuroR NS shRNA | Lentiviral transfer plasmid for expression of control shRNA | pLL3.7 PuroR | HpaI & XhoI | NS_shRNA | tgatcgacttacgacgttatttcaagagaataacgtcgtaagtcgatcttttttc | tcgagaaaaaagatcgacttacgacgttattctcttgaaataacgtcgtaagtcgatca |
| pLL3.7 PuroR JCAD shRNA | Lentiviral transfer plasmid for expression of JCAD-targeting shRNA | pLL3.7 PuroR | HpaI & XhoI | JCAD shRNA | tggtcagagcatgtgatgaattcaagagattcatcacatgctctgacctttttc | tcgaggaaaaaggtcagagcatgtgatgaatctcttgaattcatcacatgctctgacca |
| pRRL mEmerald | Lentiviral transfer plasmid for expression of JCAD fused to mEmerald at C-terminus | pRRL | EcoRI & KpnI | mEmerald | aggatcccccgggctgcaggatgtacagtgtagaagacctcc | acctccactgctgccagatctgagtccggacaccctctccactctgctag |
| pRRL JCAD-mEmerald | Lentiviral transfer plasmid for expression of JCAD fused to mEmerald at C-terminus | pRRL | EcoRI & KpnI | JCAD | aggatcccccgggctgcaggatgtacagtgtagaagacctcc | acctccactgctgccagatctgagtccggacaccctctccactctgctag |
| | | | | mEmerald | tccggactcagatctggcagcagtggaggtgtgagcaagggcgaggag | gtcattggtcttaaaggtacttacttgtacagctcgtccatg |
| pRRL mEmerald-JCAD | Lentiviral transfer plasmid for expression of JCAD fused to mEmerald at N-terminus | pRRL | EcoRI & KpnI | mEmerald | aggatcccccgggctgcaggatggtgagcaagggcgag | ctacactgtaacctccactgctgccagatc |
| | | | | JCAD | cagtggaggttacagtgtagaagacctcctg | gtcattggtcttaaaggtactcacaccctctccactctg |
| pCW57 JCAD-mEmerald | Lentiviral transfer plasmid for doxycycline-inducible expression of JCAD fused to mEmerald at C-terminus | pCW57-MCS1-P2A-MCS2 | BamHI & EcoRI | JCAD-mEmerald | gcctggagaattggctagcgatgtacagtgtagaagacc | aaggcgcaaccccaaccccgttacttgtacagctcgtc |
| pRRL mEos2-JCAD | Lentiviral transfer plasmid for expression of JCAD fused to mEos2 at N-terminus | pRRL | EcoRI & KpnI | mEos2 | aggatcccccgggctgcaggcgctagcgctaccggtcgcc | ctacactgtaaccgctgccacccccgct |
| | | | | JCAD | tggcagcggttacagtgtagaagacctcctg | gtcattggtcttaaaggtactcacaccctctccactctg |
| pRRL JCAD-HA | Lentiviral transfer plasmid for expression of JCAD fused to HA-tag at C-terminus | pRRL | EcoRI & KpnI | JCAD | aggatcccccgggctgcaggatgtacagtgtagaagacctcctg | catacgggtaacctccactgctgccagatc |

| | | | | | | |
|---|---|---|---|---|---|---|
| | | | | HA-tag | cagtggaggttacccgt atgatgttccg | gtcattggtcttaaaggt acctagctagctgcgta gtctgg |
| pRRL HA-JCAD | Lentiviral transfer plasmid for expression of JCAD fused to HA-tag at N-terminus | pRRL | EcoRI & KpnI | HA-tag | aggatcccccgggctg caggatgtacccgtatg atgttccg | tgagtccggagctagct gcgtagtctgg |
| | | | | JCAD | cgcagctagctccgga ctcagatctggc | gtcattggtcttaaaggt actcacaccctctccact ctg |
| pRRL HA-JCAD(1-450) | Lentiviral transfer plasmid for expression of JCAD(1-450) fused to HA-tag at N-terminus | pRRL | EcoRI & KpnI | HA-JCAD(1-450) | aggatcccccgggctg caggatgtacccgtatg atgttc | gtcattggtcttaaaggt acctaatcgtcaagcttt atgtc |
| pRRL HA-JCAD(1-700) | Lentiviral transfer plasmid for expression of JCAD(1-700) fused to HA-tag at N-terminus | pRRL | EcoRI & KpnI | HA-JCAD(1-700) | aggatcccccgggctg caggatgtacccgtatg atgttccgg | gtcattggtcttaaaggt acctattggggctcctcg gagaaac |
| pRRL HA-JCAD(450-700) | Lentiviral transfer plasmid for expression of JCAD(450-700) fused to HA-tag at N-terminus | pRRL HA-JCAD(1-700) | fwd: TGAGTACCTTTAAGACCAA TGACTTACAAG rev: ACCTCCACTGCTGCCAGAT C | JCAD(450-700) | gatctggcagcagtgg aggtgataaatcatata actccagtcctgtcactg | attggtcttaaaggtact cattggggctcctcgga gaaac |
| pRRL mScarlet-ZO1 | Lentiviral transfer plasmid for expression of ZO-1 fused to mScarlet at N-terminus | pRRL | EcoRI & KpnI | mScarlet | aggatcccccgggctg caggatggtgagcaag ggcgag | ctctggcggacactcga gatctgagtccg |
| | | | | ZO1 | atctcgagtgtccgcca gagctgcggcc | gtcattggtcttaaaggt acttaaaagtggtcaat aaggacagaaacaca gtttgctccaac |
| pRRL lifeact-mScarlet-P2A-HygroR | Lentiviral transfer plasmid for polycisctronic expression of lifeAct fused to mScarlet at N-terminus and hygromycin resistance casette | pRRL | EcoRI & KpnI | lifeAct | aggatcccccgggctg caggatgggtgtcgca gatttgatcaag | ccttgctcacggtggcg accggtggatc |
| | | | | mScarlet | ggtcgccaccgtgagc aagggcgaggca | cgccggatcccttgtac agctcgtccatgcc |
| | | | | P2A-HygroR | gctgtacaagggatccg gcgcaacaaac | gtcattggtcttaaaggt acctattcctttgccctcg gac |
| pLL3.7 JCAD shRNA PuroR-IRES2-JCAD-mEmerald | Lentiviral transfer plasmid for U6 promoter driven shRNA expression with polycistronic expression of puromycin resistance and JCAD-mEmerald. Silent mutations introduced into JCAD to evade shRNA knockdown. JCAD shRNA ligated into HpaI and XhoI sites as in pLL3.7 PuroR JCAD shRNA | pLL3.7 PuroR | EcoRI | IRES2 | ccgcaagcccggtgcc tgagatttaaataattcg cccctctccc | cactgtacatggtcatgg tagctccggatc |
| | | | | JCAD (fragment 1) | taccatgaccatgtaca gtgtagaagacctcc | tcttcattacatgctcgct ccttcctcc |
| | | | | JCAD (fragment 2)-mEmerald | gagcgagcatgtaatg aagaagccagtttg | agttattaggtccctcga cgttacttgtacagctcgt c |
| pLL3.7 JCAD shRNA 2 PuroR-IRES2-mEmerald | Lentiviral transfer plasmid for U6 promoter driven shRNA expression with polycistronic expression of puromycin resistance and mEmerald. | pLL3.7 JCAD shRNA PuroR-IRES2- | fwd: GGCATGGACGAGCTGTAC rev: CATGGTCATGGTAGCTCC | mEmerald | ccggagctaccatgac catggtgagcaagggc gaggag | ttgtacagctcgtccatg ccgagagtgatcccgg cggc |

| | | | | | | |
|---|---|---|---|---|---|---|
| | | JCAD-mEmerald | | | | |
| pLL3.7 JCAD shRNA 2 PuroR-IRES2-JCAD$_{(1-450)}$-mEmerald | Lentiviral transfer plasmid for U6 promoter driven shRNA expression with polycistronic expression of puromycin resistance and JCAD(1-450)-mEmerald. Silent mutations introduced into JCAD to evade shRNA knockdown. | pLL3.7 JCAD shRNA PuroR-IRES2-JCAD-mEmerald | fwd: TCCGGACTCAGATCTGGC rev: CATGGTCATGGTAGCTCC | JCAD$_{1-450}$ | ccggagctaccatgaccatgtacagtgtagaagacctc | ctgccagatctgagtccggaatcgtcaagctttatgtc |
| pLL3.7 JCAD shRNA 2 PuroR-IRES2-JCAD$_{(1-900)}$-mEmerald | Lentiviral transfer plasmid for U6 promoter driven shRNA expression with polycistronic expression of puromycin resistance and JCAD(1-900)-mEmerald. Silent mutations introduced into JCAD to evade shRNA knockdown. | pLL3.7 JCAD shRNA PuroR-IRES2-JCAD-mEmerald | fwd: TCCGGACTCAGATCTGGC rev: CATGGTCATGGTAGCTCC | JCAD$_{1-900}$ | ccggagctaccatgaccatgtacagtgtagaagacctcctgatctc | ctgccagatctgagtccggactccggcacccacatcct |
| pLL3.7 JCAD shRNA 2 PuroR-IRES2-JCAD$_{(1-1100)}$-mEmerald | Lentiviral transfer plasmid for U6 promoter driven shRNA expression with polycistronic expression of puromycin resistance and JCAD(1-1100)-mEmerald. Silent mutations introduced into JCAD to evade shRNA knockdown. | pLL3.7 JCAD shRNA PuroR-IRES2-JCAD-mEmerald | fwd: TCCGGACTCAGATCTGGC rev: CATGGTCATGGTAGCTCC | JCAD$_{1-1100}$ | ccggagctaccatgaccatgtacagtgtagaagacctcctgatctc | ctgccagatctgagtccggacaggagggactccaccgc |
| pLL3.7 JCAD shRNA 2 PuroR-IRES2-JCAD$_{(451-1359)}$-mEmerald | Lentiviral transfer plasmid for U6 promoter driven shRNA expression with polycistronic expression of puromycin resistance and JCAD(451-1359)-mEmerald. Silent mutations introduced into JCAD to evade shRNA knockdown. | pLL3.7 JCAD shRNA PuroR-IRES2-JCAD-mEmerald | fwd: TCCGGACTCAGATCTGGC rev: CATGGTCATGGTAGCTCC | JCAD$_{451-1359}$ | ccggagctaccatgaccatgaaatcatataactccagtcctgtcac | ctgccagatctgagtccggacaccctctccactctgctag |
| pLL3.7 JCAD shRNA 2 PuroR-IRES2-JCAD$_{(901-1359)}$-mEmerald | Lentiviral transfer plasmid for U6 promoter driven shRNA expression with polycistronic expression of puromycin resistance and JCAD(900-1359)-mEmerald. Silent mutations introduced into JCAD to evade shRNA knockdown. | pLL3.7 JCAD shRNA PuroR-IRES2-JCAD-mEmerald | fwd: TCCGGACTCAGATCTGGC rev: CATGGTCATGGTAGCTCC | JCAD$_{901-1359}$ | ccggagctaccatgaccatgagccctgtgtgtaggtcg | ctgccagatctgagtccggacaccctctccactctgctag |
| pLL3.7 JCAD shRNA 2 PuroR-IRES2-JCAD$_{(1101-1359)}$-mEmerald | Lentiviral transfer plasmid for U6 promoter driven shRNA expression with polycistronic expression of puromycin resistance and JCAD(1100-1359)-mEmerald. Silent mutations introduced into JCAD to evade shRNA knockdown. | pLL3.7 JCAD shRNA PuroR-IRES2-JCAD-mEmerald | fwd: TCCGGACTCAGATCTGGC rev: CATGGTCATGGTAGCTCC | JCAD$_{1101-1359}$ | ccggagctaccatgaccatgccgggcatccggagagcg | ctgccagatctgagtccggacaccctctccactctgctaggg |
| pGEX GST-mEmerald | Plasmid for bacterial expression of mEmerald fused to GST at the N-terminus | pGEX 6P-2 | EcoRI & BamHI | mEmerald | gaagttctgttccagggggcccctgggcagcagtggaggtgtgagcaagggcgaggag | atgcggccgctcgagtcgacccgggttacttgtacagctcgtccatg |

| | | | | | | |
|---|---|---|---|---|---|---|
| pGEX GST-mEmerald-JCAD | Plasmid for bacterial expression of mEmerald-JCAD fused to GST at the N-terminus | pGEX 6P-2 | EcoRI & BamHI | mEmerald-JCAD | gaagttctgttccagggg cccctgggcagcagt ggaggtgtgagcaagg gcgaggag | atgcggccgctcgagtc gacccgggtcacaccc tctccactctg |
| pGEX GST-RhoA WT | Plasmid for bacterial expression of wildtype RhoA fused to GST at the N-terminus | pGEX 6P-2 | NotI & BamHI | RhoA WT | tctgttccaggggcccct gggagctgccatccgg aagaaac | tcgtcagtcagtcacgat gctcacaagacaaggc accc |
| pGEX GST-RhoA G17A | Plasmid for bacterial expression of RhoA G17A fused to GST at the N-terminus | pGEX 6P-2 | NotI & BamHI | RhoA G17A | tctgttccaggggcccct gggagctgccatccgg aagaaac | tcgtcagtcagtcacgat gctcacaagacaaggc accc |
| pGEX GST-RhoA T19N | Plasmid for bacterial expression of RhoA T19N fused to GST at the N-terminus | pGEX 6P-2 | NotI & BamHI | RhoA T19N | tctgttccaggggcccct gggagctgccatccgg aagaaac | tcgtcagtcagtcacgat gctcacaagacaaggc accc |
